# 3D Chromatin Architecture Remodeling during Human Cardiomyocyte Differentiation Reveals A Role Of HERV-H In Demarcating Chromatin Domains

**DOI:** 10.1101/485961

**Authors:** Yanxiao Zhang, Ting Li, Sebastian Preissl, Jonathan Grinstein, Elie N. Farah, Eugin Destici, Ah Young Lee, Sora Chee, Yunjiang Qiu, Kaiyue Ma, Zhen Ye, Rong Hu, Quan Zhu, Hui Huang, Rongxin Fang, Leqian Yu, Juan Carlos Izpisua Belmonte, Jun Wu, Sylvia M. Evans, Neil C. Chi, Bing Ren

**Author notes:** Correspondence: Neil Chi and Bing Ren. these authors contributed equally (Yanxiao Zhang, Ting Li, Sebastian Preissl).

## Abstract

Dynamic restructuring of chromatin architecture has been implicated in cell-type specific gene regulatory programs; yet, how chromatin remodels during lineage specification remains to be elucidated. Through interrogating chromatin reorganization during human cardiomyocyte differentiation, we uncover dynamic chromatin interactions between genes and distal regulatory elements harboring noncoding variants associated with adult and congenital heart diseases. Unexpectedly, we also discover a new class of human pluripotent stem cell (PSC)-specific topologically associating domains (TAD) that are created by the actively transcribed endogenous retrotransposon HERV-H. Deletion or silencing of specific HERV-H elements eliminates corresponding TAD boundaries, while *de novo* insertion of HERV-H can introduce new chromatin domain boundaries in human PSCs. Furthermore, comparative analysis of chromatin architecture in other species that lack HERV-H sequences supports a role for actively transcribed HERV-H in demarcating human PSC-specific TADs. The biological role of HERV-H is further underscored by the observation that deletion of a specific HERV-H reduces transcription of genes upstream and facilitates cell differentiation. Overall, our results highlight a previously unrecognized role for retrotransposons in restructuring genome architecture in the human genome and delineate dynamic gene regulatory networks during cardiomyocyte development that inform how non-coding genetic variants contribute to human heart diseases.

## MAIN TEXT

The three-dimensional organization of chromosomes enables long-range communications between enhancers and promoters that are critical for building complex gene regulatory networks in multicellular species^1,2^. In somatic cells, interphase chromosomes occupy separate nuclear spaces known as chromosome territories^3^. Each chromosome is organized into a dynamic but non-random hierarchical structure characterized by stretches of transcriptionally active, megabase-long compartments that are interspersed with stretches of transcriptionally inactive compartments^4^. These compartments can be further partitioned into topologically associating domains (TADs), which exhibit high levels of intra-domain interactions and relatively low levels of inter-domain interactions^5–7^. The boundaries of TADs are generally conserved between cell types and closely related species^5,8^, and can restrict *cis*-regulatory element interactions with target promoters^9^; therefore when TADs are disrupted, it can lead to significant change of transcriptional landscape and cause diseases^10–13^. At or within TAD boundaries, long-range interactions such as chromatin loops are frequently observed to link regulatory elements, including binding sites for the CCCTC-binding factor (CTCF), promoters and enhancers^14,15^.

There is mounting evidence that a large fraction of TADs and chromatin loops are shaped by cohesin mediated loop extrusion^16–18^, with TAD borders constrained by the CTCF^19,20^. However, not all TAD boundaries contain CTCF binding sites, and many CTCF binding sites do not localize on or near TAD boundaries^5^, suggesting that their underlying relationship is more complex and raising the question that additional mechanisms for TAD formation may also exist. On the other hand, CTCF binding sites are dynamically evolving and underlie the evolution of genome architecture^21^. Interestingly, a subset of species-specific CTCF binding sequences are rapidly dispersed by transposable elements, especially SINE elements^22^. This leads to an important question that has not been thoroughly investigated: how much a role do transposable elements play in reshaping the genome architecture during evolution?

Genome architecture is not only reshaped during evolution, but also reconfigured in a cell-type specific manner during stem cell differentiation and somatic cell reprogramming^8,23–26^. Failure of proper restructuring of chromatin has potentially detrimental effects and might lead to developmental defects^10^. One of the most common birth defects is congenital heart disease (CHD) which impacts 1% of newborns^27^. Mutations in chromatin modifiers, remodelers as well as transcription factors have been linked to CHD^28^. In addition, it was reported that cardiomyocyte-specific deletion of CTCF was sufficient to induce heart failure in mice^29^, highlighting the importance of chromatin architecture in cardiac development. Two recent studies showed promoter centric chromatin interaction maps of in vitro differentiated cardiomyocytes^30,31^. However, these studies did not capture the dynamics during the differentiation process.

Here, using high-resolution *in situ* high-throughput chromosome conformation capture Hi-C (*in situ* Hi-C)^14,15^, we interrogated the reorganization of hPSC chromatin architecture during key developmental stages of human cardiomyocyte differentiation. We report the finding of stage-specific long-range chromatin interactions that are associated with temporal regulation of gene expression and predict candidate genes for non-coding DNA variants associated with cardiac traits/diseases. Moreover, we reveal a class of primate-specific endogenous retrotransposon, HERV-H, that is involved in establishing boundaries of TADs *de novo* in hPSCs during primate evolution. Importantly, we show that HERV-H demarcates TAD boundaries, which are dependent on its transcription, as targeted silencing of HERV-H can weaken or eliminate the insulation of TAD borders. Finally, comparative analysis of chromatin architecture of PSC in different primate and non-primate species, as well as creation of new hPSC TADs via *de novo* insertions of HERV-H further underscore this conclusion. Taken together, our study emphasizes an important role for ERV family retrotransposon elements in shaping chromatin organization in specific cell types, and provides a valuable resource for the study of human cardiac development and disease.

## RESULTS

### Dynamic chromatin organization during cardiomyocyte differentiation

To study the dynamics of genome organization during cardiomyocyte development, we utilized a transgenic human embryonic stem cell (hESC) H9 *MYL2:H2B-GFP* ventricular cardiomyocyte reporter line, which can be used to purify ventricular cardiomyocytes^32^ (Extended Data Fig. 1a). We generated and collected samples in biological duplicates at the following six critical time points of cardiomyocyte differentiation: hESCs (Day 0), mesodermal cells (Day 2), cardiac mesodermal cells (Day 5), cardiac progenitors (Day 7), primitive cardiomyocytes (Day 15) and ventricular cardiomyocytes (Day 80) (Fig. 1a). Flow cytometry confirmed that differentiation efficiency for D2 – D15 time point samples was at least ~80-90%, and *MYL2:H2B-GFP*+ ventricular cardiomyocytes were sorted at day 80 (Extended Data Fig. 1b). *In situ* Hi-C^4^ experiments were performed, and on average three billion raw read pairs and one billion unique long-range cis contacts (minimum distance 10 kb, Extended Data Table 1) were generated for each time point. Close correlation of Hi-C maps between biological replicates verified that these maps were highly reproducible at multiple scales (Extended Data Fig. 2). Complementing these Hi-C contact maps, ATAC-seq, ChIP-seq and RNA-seq experiments were performed on these samples to interrogate chromatin accessibility, identify CTCF occupancy sites and key histone modifications, and profile transcriptomes, respectively (Extended Data Fig. 3).

**Figure 1.**
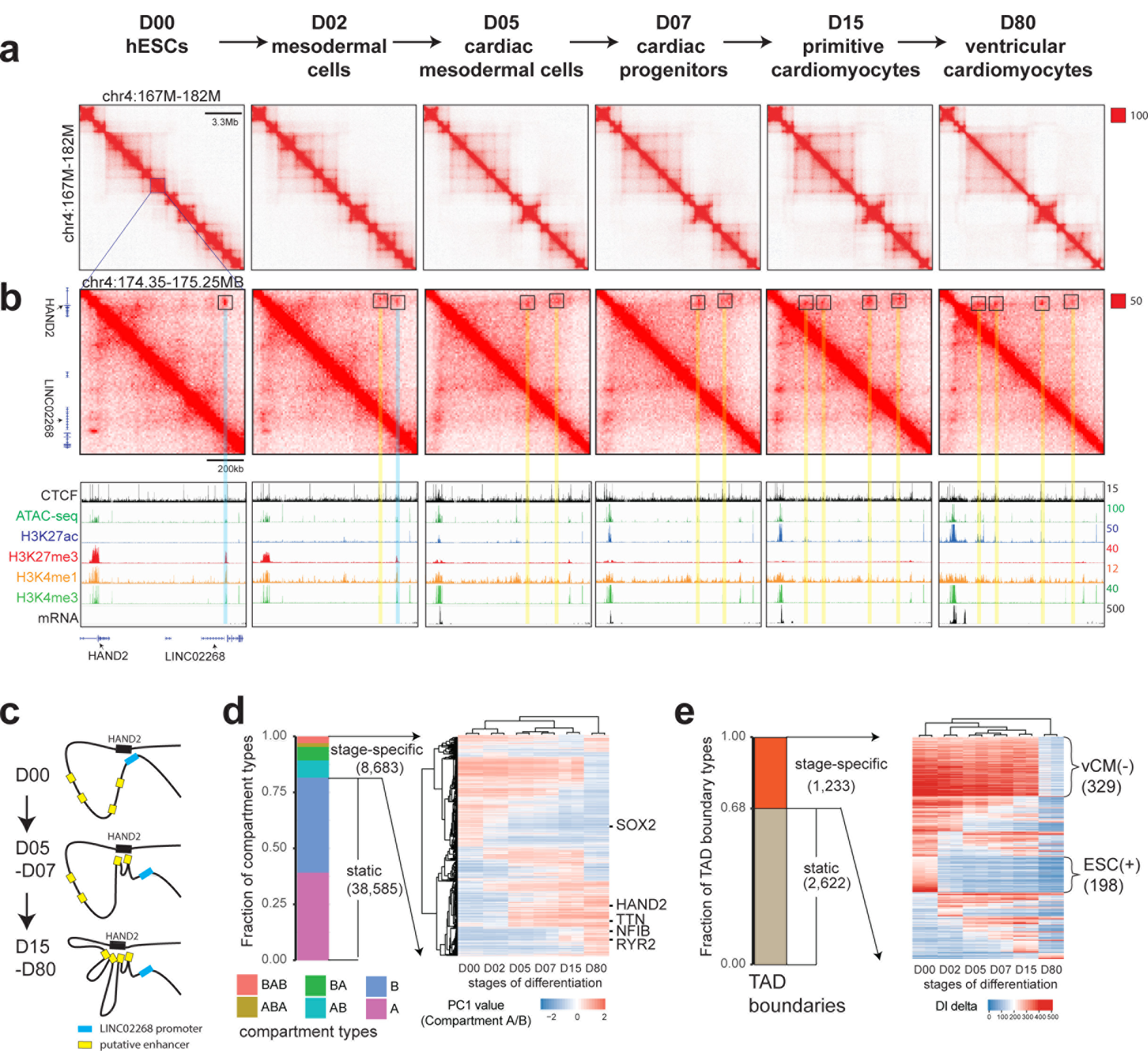
Global chromatin architecture reorganization during human ventricular cardiomyocyte differentiation. (**a & b**) Hi-C contact matrices for each stage of cardiomyocyte differentiation at mega-base resolution and at higher resolution at the *HAND2* locus with corresponding epigenomic profiles. (**c**) Schematic illustration of the multi-step decommissioning of inhibitory loop and assembly of subsequent activating loops anchored at *HAND2* promoter. (**d**) A heatmap showing hierarchical clustering of dynamic chromatin compartments during cardiomyocyte differentiation. The pseudo color reflects the PC1 values (compartment A/B) of compartment bins. Negative PC1 value stands for compartment B and positive for compartment A. Representative genes located in corresponding compartment bins are annotated to the right of the heatmap. A heatmap showing the DI delta scores for the stage-specific TAD boundaries, ordered by the presence of TADs at six stages. vCM(-) stands for TAD boundaries lost in purified ventricular cardiomyocytes at D80, ESC(+) for hESC-specific TAD boundaries.

Analyses of these genome-wide datasets revealed extensive reorganization of chromatin architecture at all structural levels during hESC differentiation, which was also accompanied by corresponding epigenomic changes (Fig. 1a, b, c). Across the entire genome, the abundance of short-range chromatin interactions decreased while the number of long-range chromatin interactions (above 500 kilobases) increased (Extended Data Fig. 4a). Identification of active (A) and inactive (B) compartments at all stages using principal component analysis showed that 81.6% of the genome maintained the same compartment state throughout cardiomyocyte differentiation (Fig. 1d). Among the 18.4% of the genome that switched compartments, a similar proportion of the genome switched from A to B and from B to A at each stage transition (Extended Data Fig. 4b), and the majority (73%) only switched once, suggesting linear nuclear architectural changes during lineage commitment (Fig. 1d). Hierarchical clustering of genomic bins that switched compartments showed stage specific dynamics (Fig. 1d), which positively correlated with gene expression (Extended Data Fig. 4c) as reported previously^8^. For instance, gene loci of pluripotency factors including *SOX2* moved to the inactive compartment when hESCs differentiated into mesodermal cells, whereas genes involved in cardiomyocyte specification such as the transcription factor *HAND2* and the cardiomyocyte-specific calcium channel *RYR2* displayed B to A transitions at day 5 and day 15, respectively (Fig. 1d; Extended Data Fig. 4d).

### Transcription of HERV-H is associated with TAD boundary formation in human ES cells

Consistent with recent reports describing loss of TADs during the differentiation of embryonic stem cells^23,33^, we also observed that the number of TADs decreased as hESCs differentiated into ventricular cardiomyocytes using multiple TAD-calling algorithms^5,14,34^ (Extended Data Fig. 5a, b, c). Clustering of stage-specific TAD boundaries using Directionality Index (DI) delta scores, which estimates the strength of the TAD boundaries, further showed that the majority of stage-specific TAD boundaries were lost or weakened during differentiation and the effect was most pronounced at early mesoderm specification (D0-D2), and ventricular cardiomyocyte differentiation/maturation (D15-D80) (Fig. 1e). To investigate the mechanism of TAD dynamics during cardiomyocyte differentiation, we examined the sequence features of TAD boundaries that were lost in the course of differentiation with a particular focus on D0 hESC-specific TAD boundaries [ESC(+)] and the boundaries lost during D15-D80 ventricular cardiomyocyte transition [vCM(-)] (Extended Data Table 2). While CTCF has been shown to play a critical role in defining TAD boundaries^5,14,20^, we observed no significant difference in CTCF occupancy across all stages examined at ESC(+), vCM(-) and stable TAD boundaries (Extended Data Fig. 5d). Thus, we explored whether other potential mechanisms might be involved in the formation of these TAD boundaries. Interestingly, we discovered that eight classes of repeat elements were over-represented specifically at ESC(+) TAD boundaries when compared to stable TAD boundaries (p-value < 0.01 and fold change > 2), whereas no enrichment of these repeats was observed in vCM(-) TAD boundaries (Extended Data Fig. 6a). Additionally, the ESC(+) TAD boundaries were associated with hESC-specific H3K27ac signal and mRNA transcription (Extended Data Fig. 5e, f), which was particularly enriched for HERV-H transcription (Fig. 2a and Extended Data Fig. 6b).

**Figure 2.**
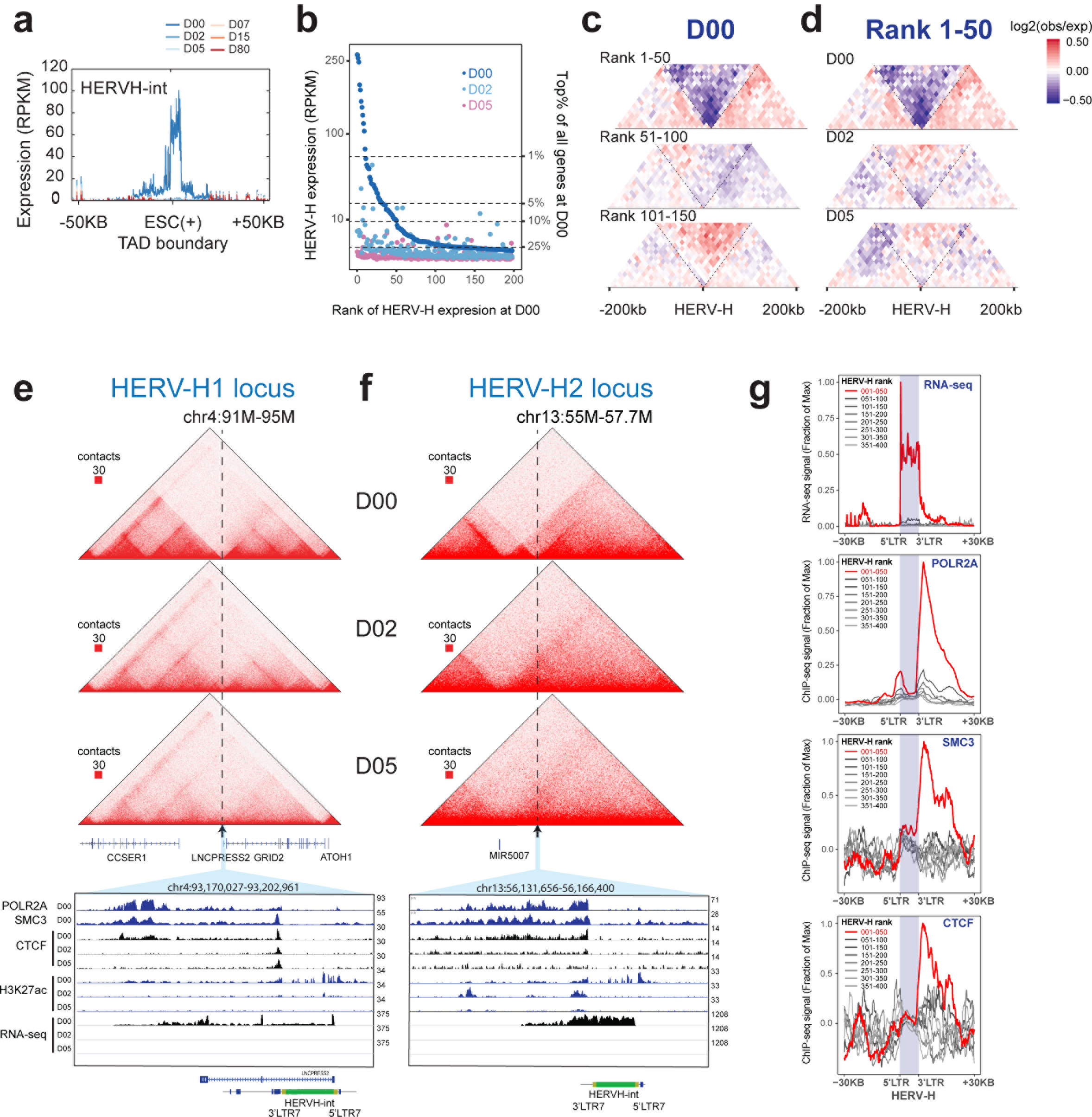
Transcriptionally active HERV-H forms human ESC-specific TAD boundaries. (**a**) Aggregate RNA-seq expression profile (RPKM normalized) at ESC(+) TAD boundaries that overlap HERV-H element. (**b**) Scatterplot shows the expression levels (RPKM) across different HERV-H loci (ordered by expression levels from high to low in ESC) at D0, D2 and D5 stages of differentiation. (**c**) Heatmap of aggregate Hi-C contact matrix [log2(observed/expected)] within 200 kb of the top 50, 51-100 and 101-150 ranked HERV-Hs, at D0. (**d**) Heatmap of the aggregated Hi-C matrix [log2(observed/expected)] within 200 kb of the top 50 HERV-Hs, at D0, D2 and D5. (**e,f**) Representative Hi-C interaction matrices of two HERV-H loci located at ESC(+) TAD boundaries at D0, D2 and D5 (top) are shown as heatmaps along with genome browser tracks of POLR2A, SMC3, CTCF, H3K27ac ChIP-seq and RNA-seq data of the expanded genomic region containing the TAD boundary (arrow). (**g**) Aggregated genomic profiles of RNA-seq, POLR2A, SMC3 and CTCF ChIP-seq around top 50 HERV-Hs located on the ESC(+) TAD boundaries (red) and lower ranked HERV-Hs (grey).

HERV-H is a class of primate-specific endogenous retrotransposons. It is transcriptionally active in both human preimplantation embryos and pluripotent stem cells^35,36^, and plays a critical role in pluripotency, stem cell maintenance and somatic cell reprograming^37–41^. Although more than 1000 copies of HERV-H sequences exist in the human genome and are expressed at different levels in hESCs (Fig. 2b)^37^, we discovered that the top 50 highly transcribed HERV-H loci were predominantly located at TAD boundaries in hESCs (Fig. 2c, Extended Data. Fig 6c). Notably, transcriptional silencing of these HERV-Hs was observed by day 2 in hESCs differentiating into mesodermal cells and coincided with the loss of the corresponding TAD boundaries (Fig. 2d; specific loci in Fig2e, f and Extended Data Fig 6d, e, f). As controls, genes with similar transcriptional levels to these top 50 HERV-H loci did not display any boundary activity as evidenced by the DI scores (Extended Data Fig. 7a). This correlation was present in not only H9 hESCs but also other hPSCs including H1 hESCs^8,42^ and human induced PSCs (iPSC)^43^; however, differentiated cells did not exhibit these HERV-H associated TAD boundaries (Extended Data Fig. 7b).

Because HERV-Hs might contribute to TAD boundary formation by recruiting DNA binding proteins, we examined publicly available ChIP-seq data of histone modifications and transcription factors (TFs) in hESCs. We identified several modifications and factors enriched at sites for the top 50 transcribed HERV-Hs when compared to those for the lower-ranking HERV-Hs. They included POL2 transcriptional machinery (POLR2A and POLR2AphosphoS5), active histone marks (H3K4me3, H3K4me2 and H3K27ac), ubiquitously active TFs (JUND and SP1), pluripotency factor NANOG^39^ and all three subunits of the cohesin complex (Extended Data Fig. 7c). To further investigate the role of cohesin in HERV-H-mediated TAD boundary formation, we performed ChIP-seq in hESCs for the cohesin complex subunit SMC3. Analyzing this ChIP-seq data along with RNA-seq as well as CTCF and POLR2A ChIP-seq data, revealed broad enrichment of cohesin, CTCF and POLR2A specifically at the 3’ end of these highly transcribed HERV-H sequences (Fig. 2g). In contrast to these findings, the enrichment of these factors was not similarly observed at sites of lower ranked HERV-Hs (Fig. 2g), suggesting that these factors may be involved in mediating the formation of TAD boundaries by HERV-Hs. However, we cannot completely rule out that CTCF may also participate in defining these HERV-H-associated TAD boundaries, as weak but reproducible enrichment of CTCF occupancy was observed in these regions despite the absence of CTCF binding motifs (Fig. 2g). Overall, these data support that HERV-Hs form TAD boundaries by accumulating cohesin complexes at its 3’ end, and cohesin is likely positioned by the transcribing POL2 complex^44,45^.

### Deletion or silencing of HERV-H elements abolishes the corresponding TAD boundaries

To determine whether HERV-H sequences are indeed required for TAD boundary formation, we deleted two HERV-H elements located at two of the strongest hESC TAD boundaries, using CRISPR-Cas9 genome editing tools (termed as HERV-H1-KO and HERV-H2-KO, respectively). In both cases, deletion of these HERV-Hs resulted in elimination of TAD boundaries, as evidenced by the merging of the two TADs bordering each HERV-H element (Fig. 3a), and reduced expression of genes in the TAD domain immediately 5’ upstream of the HERV-H sequence (Fig. 3a). These findings thus reveal that HERV-H sequences are indispensable for the TAD boundaries formed at these regions and furthermore the presence of HERV-H increases the overall transcriptional activity of the neighboring genes in the same TAD domain, supporting that HERV-Hs may serve as enhancers in hESCs^46,47^. Notably, this activating effect is only seen in the TAD domain on the 5’ end of HERV-H and not the 3’ end, suggesting that boundaries present on the 3’ end of HERV-Hs, may insulate the effect of enhancers. Supporting this possibility, additional analyses of all genes located within 500 kb of boundary-associated HERV-Hs showed that only the genes in regions 5’ of HERV-H were up-regulated in hESCs compared to differentiated cells (Fig. 3b).

**Figure 3.**
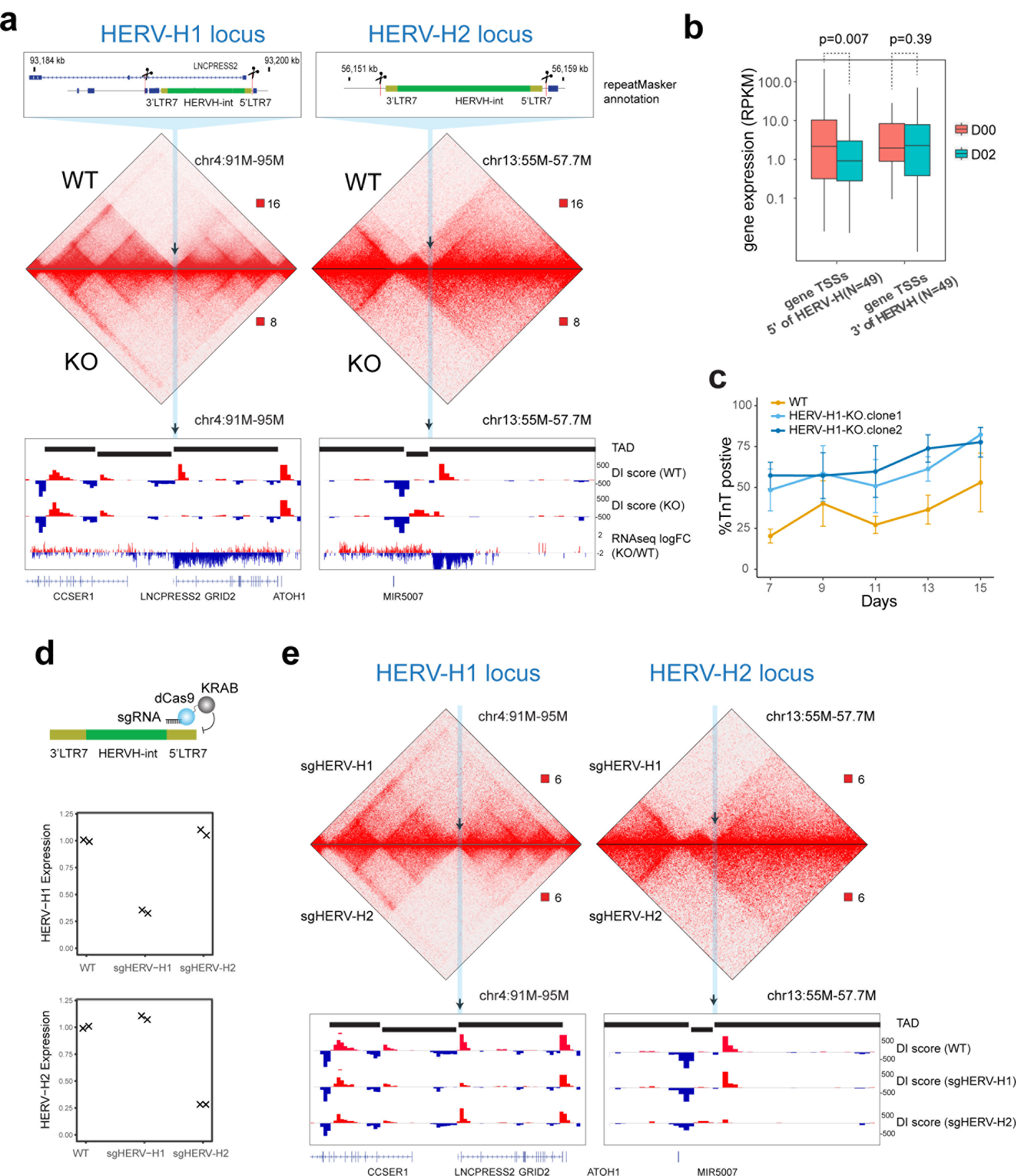
Deletion or silencing of two HERV-H sequences leads to merging of TADs in hESC. (**a**) Hi-C interaction matrices of the wild-type (WT) and mutant hESC lines (HERV-H1-KO and HERV-H2-KO) are shown, along with DI scores, and fold changes of gene expression at the HERV-H1 and HERV-H2 loci. The loss of TAD boundary in the mutant cells is accompanied with decrease of RNA expression 5’ terminus to the HERV-H sequences. (**b**) Boxplots show expression levels (RPKMs) of genes whose TSSs are located from −500 kb to the 5’ LTR and from 3’ LTR to +500 kb of HERV-Hs of boundary-associated HERV-Hs. (**c**) Line chart (mean ± standard error, N = 3 cardiomyocyte differentiations) shows percentage of TNNT2 positive cells during cardiomyocyte differentiation of WT and HERV-H1-KO hESC lines (two HERV-H1-KO clones analyzed). HERV-H1-KO hESCs display increased cardiomyocyte differentiation efficiency compared to WT hESCs. (**d**) Design of the CRISPR-dCas9-KRAB system to silence HERV-H expression (top) and gene expression values of HERV-H1 and HERV-H2 in WT and CRISPR-i targeted hESCs (bottom). (**e**) Hi-C interaction matrices of the CRISPR-i targeted hESC lines (sgHERV-H1 and sgHERV-H2) are shown, along with DI scores at the HERV-H1 and HERV-H2 loci.

Further examination of the transcriptional profiles of both HERV-H knockouts revealed that expression of 353 and 144 genes were significantly changed (Extended Data Fig. 8a, b, Extended Data Table 3), and the fold changes of HERV-H1-KO and HERV-H2-KO were highly concordant (Extended Data Fig. 8c, Pearson r=0.5, p-value< 1e-15) despite both engineered cells maintaining pluripotency. Of the 43 genes down-regulated in both KOs, 10 (23%) were located within 20 kb of other HERV-H sequences (Extended Data Fig. 8d), including two well studied chimeric transcripts *SCGB3A2*^48^ and *LINC00458*^49^, which are known to be regulated by HERV-Hs (Extended Data Fig. 8e, f). Thus, these data suggest that deletion of individual HERV-H sequences could lead to down-regulation of HERV-H related transcription at other loci. In addition to these findings, we also discovered that the human-specific long-coding RNA (lncRNA), *Heart Brake* (*HBL1*), which suppresses cardiomyocyte differentiation^50^, was also significantly reduced in the HERV-H1-KO (Extended Data Fig. 8f). To further explore whether these changes in transcriptional landscape in the HERV-H1-KO promotes cardiomyocyte differentiation, we examined the cardiomyocyte differentiation efficiency of HERV-H1-KO hESCs compared to control hESCs under suboptimal cardiomyocyte differentiation conditions. While these conditions led to ~40% cardiomyocyte differentiation efficiency at day 15 in control hESCs, we discovered that HERV-H1-KO hESCs exhibited >80% cardiomyocyte differentiation efficiency (Fig. 3c). Altogether, these results suggest that removal of one highly expressed TAD-associated HERV-H sequence might facilitate the exit from pluripotency, including the cardiac lineage examined in this study.

Having established that HERV-H DNA sequences are required for TAD boundary formation, we investigated whether its transcriptional activity is also essential. To address this question, we blocked the transcription of the aforementioned HERV-H loci without altering the HERV-H DNA sequence by employing a CRISPR-dCas9-KRAB system with sgRNAs that targeted the promoters (5’LTR7) of these HERV-Hs (Fig. 3d). Each sgRNA specifically reduced the targeted HERV-H expression by over 70%, while not affecting the expression of other HERV-Hs (Fig. 3d). Accompanying the reduction of HERV-H, the TAD border was greatly weakened, evidenced by the decreased directionality index (Fig. 3e). This result indicates that HERV-H’s high transcription is required for creating these groups of TAD boundaries in hESC.

### HERV-H introduces *de novo* TAD boundaries

It is estimated that HERV-H integrated into the primate lineage 30–40 million years ago (MYA) at the time of divergence of Old and New World monkeys^51,52^, with the largest expansion (LTR7 and LTR7B) occurring in the Old World monkey lineage, including the Great apes^51^. Consequently, the Great Apes, including humans, harbors over 1000 copies of HERV-H, whereas New World monkey has 50-100 copies and none in non-primates. We hypothesize that HERV-H might have introduced *de novo* chromatin domains during primate evolution. To test this hypothesis, we performed Hi-C to interrogate the chromatin architecture of chimpanzee iPSC, marmoset (New World monkey) iPSC and mouse ESC, and compared them to that of the human ESC. While the top 50 transcribing HERV-H loci showed contact insulation in human ESC, their syntenic regions in the marmoset iPSC and mouse ESC did not exhibit such insulation (Fig. 4a). Surprisingly, although chimpanzee has similar copies of HERV-Hs as human, we did not observe any TAD boundaries in the syntenic regions (Fig. 4a). One reason for the lack of boundaries in chimpanzee could be the significantly lower expression of HERV-Hs in chimpanzee iPSC under our culturing conditions (Extended Data. Fig. 9a). Taken together, our data suggests that HERV-H introduces *de novo* TAD boundaries in PSCs during primate evolution, in a transcription-dependent manner. Furthermore, additional sequence analysis of the HERV-H LTR sequence revealed that the TAD-forming HERV-Hs were predominantly flanked by a 450bp subtype of LTR7 at both ends (Extended Data Fig. 9b, c). And those HERV-Hs showed less sequence divergence between their 5’ LTRs and 3’ LTRs (Extended Data Fig. 9d), suggesting that they likely were inserted more recently than other HERV-Hs that do not form TAD boundaries, or were impacted by gene conversion or selection^53^.

**Figure 4.**
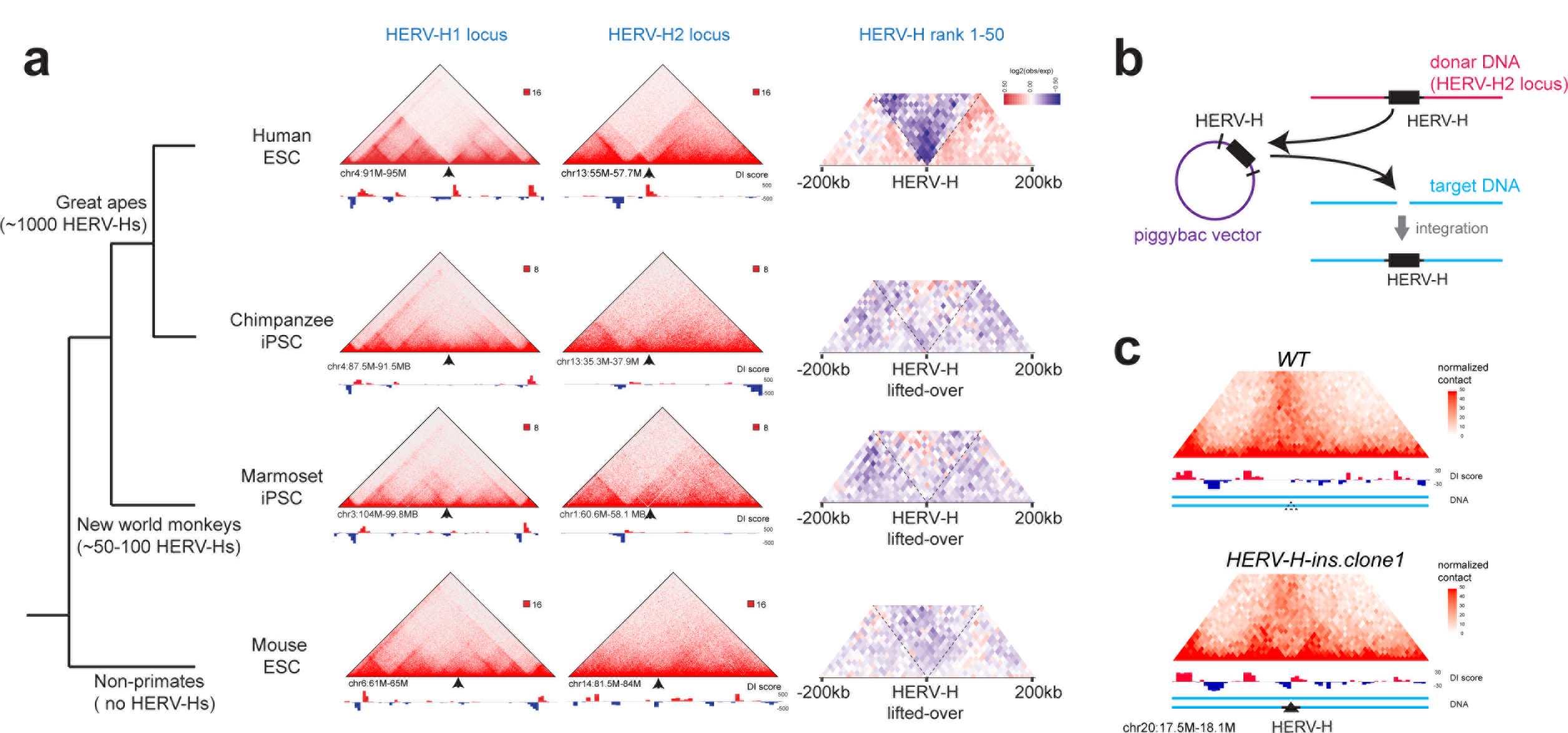
HERV-H introduces *de novo* TAD boundaries during primate evolution and in engineered hESC. (**a**) Simplified tree of primate evolution with the copies of HERV-H annotated (left) and Hi-C interaction matrices of human ESC, chimpanzee iPSC, marmoset iPSC and mouse ESC are shown, along with DI scores at the syntetic regions to human HERV-H1 locus, HERV-H2 locus, and all top 50 transcribing HERV-Hs. (**b**) Design of the piggybac vector to “transpose” HERV-H to random genomic locations in the hESC (HERV-H2-KO line). (**c**) Hi-C interaction matrices of the parental cell line (HERV-H2-KO) and the cell line with HERV-H insertions (HERV-H-ins.clone1) are shown, along with DI scores at the locus of one HERV-H insertion.

Finally, to investigate whether the *de novo* insertion of HERV-Hs can create new TAD boundaries, we engineered two human hESC lines in which an 8kb sequence from the HERV-H2 locus (with 1kb flanking HERV-H from each side) was cloned and randomly inserted it into the H9 hESC genome using a piggybac transposon (Fig. 4b). From sequence analyses, we discovered that this HERV-H2 construct was inserted into 52 new genomic loci (Extended Data Fig. 9e,f), of which six exhibited a significant increase in the local contact insulation at the place of insertion, as evidenced by the change in Hi-C contact matrix and DI score (One example in Fig. 4c). This data demonstrated that HERV-H insertion can create chromatin boundaries *de novo*, however, such ability is likely also influenced by the native genomic contexts.

### Stage-specific chromatin loops form interaction networks around cardiac transcription factors

From our extensive interrogation of the genome during hESC cardiomyocyte differentiation, we also identified 14,216 chromatin loops spanning a median genomic distance of 180 kb at 10 kb resolution (Extended Data Table 4). More than 70% of the loops corresponded to either promoter-to-promoter or promoter-to-enhancer interactions (Extended Data Fig. 10a). Consistent with chromatin loops detected in different cell types or model systems^14,15^, the loop anchors were enriched for accessible chromatin, histone marks, transcriptional start sites and CTCF binding sites (Extended Data Fig. 10b). More importantly, 23% (3,251) of these chromatin loops exhibited significant changes in interaction strength during cardiomyocyte differentiation, which illustrates a dramatic reconfiguration of the gene regulatory network (Fig. 5a, Extended Data Fig. 10c). Gene transcription and chromatin marks at anchors of stage-specific chromatin loops were correlated to a significantly higher degree than that of anchors of static chromatin loops and distance-matched control (Fig. 5e). Using K-means clustering, we defined five distinct clusters of stage-specific chromatin loops (Fig. 5b), each accompanied by corresponding epigenomic changes (Extended Data Figure 10d). Remarkably, the gene sets enriched for each chromatin loop cluster recapitulated the developmental processes from early differentiation, to heart development, and culminated in cardiac muscle development (Fig. 5c). Chromatin loops gained during cardiomyocyte differentiation included promoters of many well studied heart transcription factors, such as *GATA6*, *TBX5*, *NKX2-5*, *TBX3* and *MEF2A*^54^. Analysis of ATAC-seq peaks at loop anchors also revealed distinct patterns of transcription factor binding throughout differentiation (Fig. 5d). For instance, enrichment for GATA motifs emerged at day 2, and T-BOX motifs at day 5, whereas MEF2 motifs were only highly enriched at day 80. These findings suggest a process involving pioneering factor activity of GATA family members such as GATA4 for the initial rewiring of chromatin loops to activate relevant early cardiac developmental gene programs, whereas MEF family members such as MEF2C may play a role in the maturation of cardiomyocytes. Finally, loops formed at later stages tended to span longer genomic distances, to have a higher percentage of convergent CTCF-CTCF motifs, and were more likely to occur at TAD boundaries (Extended Data Fig. 10e, f, g). These observations further support the global trend of increased long-distance interactions during cardiomyocyte differentiation (Extended Data Fig. 4a).

**Figure 5.**
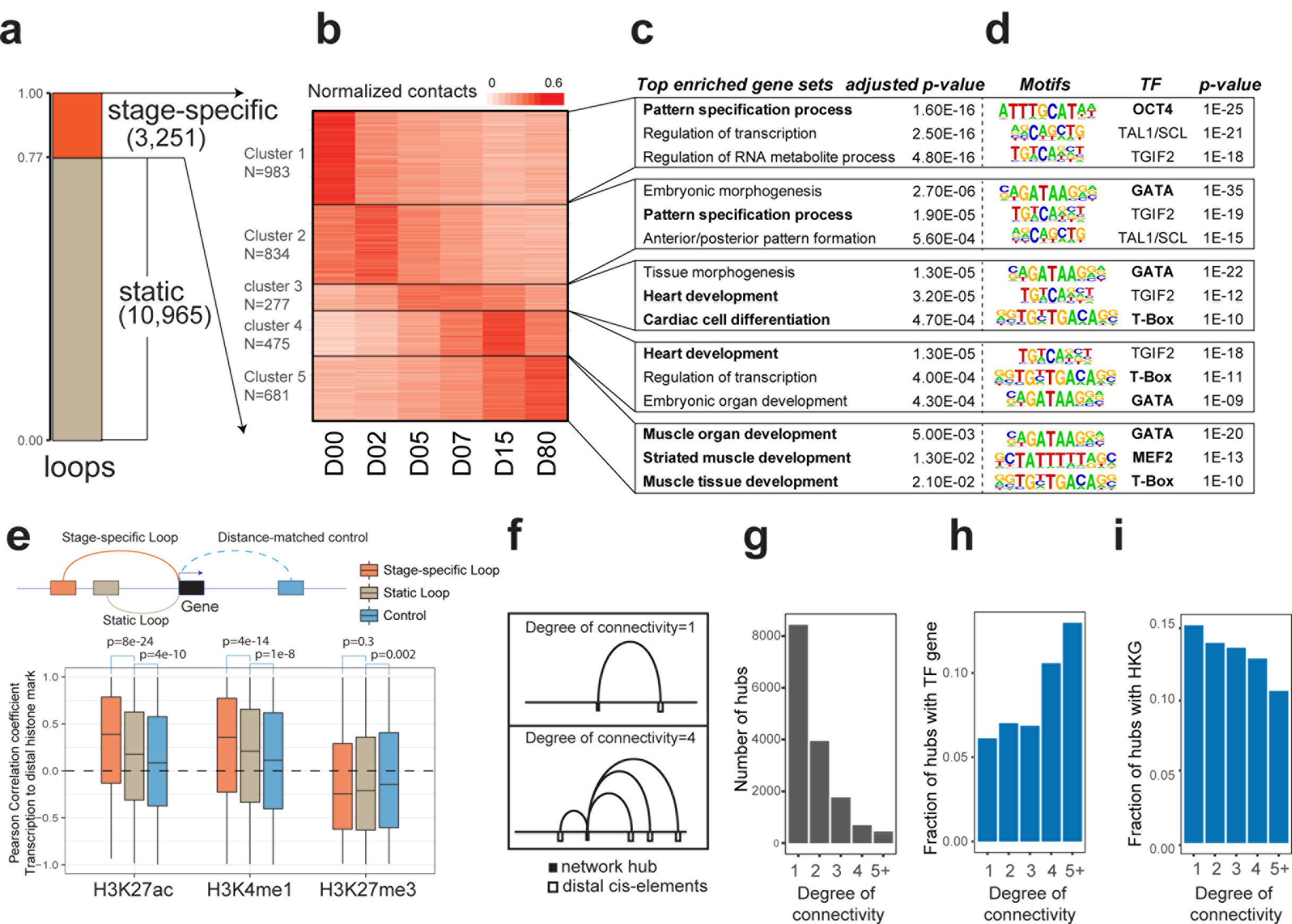
Dynamics of chromatin loops during human cardiomyocyte differentiation. (**a**) Bar-chart shows the percentages of static or dynamic chromatin loops during cardiomyocyte differentiation. (**b**) Heatmap shows K-means clustering of the stage-specific chromatin loops. Pseudo color reflects the normalized contact frequencies between the loop anchors for each stage-specific loop. (**c**) Enriched Gene Ontology (GO) terms for genes located at the loop anchors of each cluster in (**b**) along with their adjusted p-values. (**d**) Over-represented transcription factor binding motifs for distal open chromatin regions located at the loop anchors of each cluster in (**b**) and their p-values. CTCF motif enrichment was not shown but observed in all clusters. (**e**) Boxplots of Pearson correlation coefficients of histone marks of distal enhancers and expression levels of their target genes predicted based on the chromatin loops. Distance-matched controls are “mirror” loops that connect to genomic regions from the opposite direction with the same distances. P-values are from Wilcoxon Rank Sum test. (f) Diagram illustrates how the degree of chromatin interactions for each loop anchor is determined. (**g**) Histogram of degrees of network connections for all loop anchors. (**h**) Fraction of chromatin interaction network hubs containing at least one transcription start site (TSS) of a transcription factor (TF) plotted against the degree of connectivity. (**i**) Fraction of network hubs containing at least one TSS of a house-keeping gene (HKG) plotted against the degree of connectivity.

Integrative analysis of the chromatin architecture and ChIP-seq data revealed two major types of loops: 1) loops enriched for the polycomb mark H3K27me3 (Extended Data Fig 10d, 11a); and 2) loops enriched for H3K27ac (Extended Data Fig 10d, 11b). H3K27me3-associated, polycomb dependent loop interactions, which have been described in *Drosophila* and mouse ESC^23,55,56^, were prominent in hESCs, spanned very long genomic distances and were abruptly lost at the mesoderm stage (Extended Data Fig 10d, 11a). Notably, such loops frequently involved promoters of transcription factors, which were not expressed at the hESC stage (Fold enrichment=9.7, adjusted p-value=1e-70, hypergeometric test), including the early cardiac transcription factor *HAND2*. Further analyses of the *HAND2* locus revealed a multistep assembly of loops formed between the *HAND2* promoter and several distal elements (Fig. 1b). In hESCs and mesodermal cells, which do not express *HAND2*, the *HAND2* promoter formed an H3K27me3 associated loop with the promoter region of the lncRNA *LINC02268*. However, by day 5, this loop was eliminated and two new loop interactions were established between the *HAND2* promoter and two distal regions marked by H3K27ac and H3K4me1. This chromatin remodeling coincided with the onset of *HAND2* expression in cardiac mesodermal cells (D5) and a subsequent 4-fold increase in *HAND2* expression in primitive cardiomyocytes (D15) and ventricular cardiomyocytes (D80); however, lncRNA *LINC02268* remained transcriptionally repressed (Fig. 1b, c). Based on this multi-loop chromatin remodeling at the *HAND2* locus, we further examined whether multi-loop formation was a general feature to tightly control expression levels of transcription factors whose function may impact cell fate more than that of other genes. To this end, we calculated the degree of interaction for each loop anchor (Fig. 5f), and observed that 45% of the loop anchors interacted with at least two other anchors (Fig. 5g), suggesting that chromatin loops frequently form network hubs. In addition to *HAND2*, many other well-known cardiac TFs, including *TBX5*, *GATA4*, *TBX20*, *MEIS1/2*, also interacted with four or more *cis*-elements and formed similar network hubs (Extended Data Fig 11c, d, e, and Extended Data Table. 5). Supporting these findings, the network hubs with more chromatin interactions were more likely to include transcription start sites (TSSs) of TFs (Fig. 5h), and less likely to involve TSSs of house-keeping genes (Fig. 5i). Finally, those network hubs with more interactions frequently contained more CTCF peaks (Extended Data. Fig. 11f), suggesting that these interactions were likely facilitated by CTCF.

### Chromatin loops predict cognate target genes for cardiac GWAS SNPs

We next used the chromatin interaction maps to link non-coding genetic variants from genome-wide association studies (GWAS) to potential target genes, and found a strong enrichment of single nucleotide polymorphisms (SNPs) associated with cardiac traits in the chromatin loop anchors (Fig. 6a). The list of predicted candidate genes for these SNPs predominantly included genes involved in cardiac function (Extended Data Table 6) and cardiac ion channel genes (Fig. 6b). One notable cardiac GWAS example was rs6781009, which was found in strong linkage disequilibrium (LD) with lead SNPs of four cardiac electrophysiologic GWAS traits (QRS complex, QRS duration, QT interval and electrocardiographic traits). Using our chromatin interaction maps, we found that this SNP was located in a stage-specific D80 enhancer that interacted with the promoter of the cardiac sodium channel gene *SCN5A* (Extended Data Fig. 12a), supporting similar findings recently observed by circular chromosome conformation capture (4C) studies in human cardiac tissue and functional studies in mouse hearts^57^. Another cardiac GWAS example was rs17608766, the lead SNP for six cardiac traits including aortic root size, blood pressure, pulse pressure, QRS complex, QRS duration and systolic blood pressure. Our chromatin interaction maps revealed that this SNP overlapped a D2 mesoderm specific enhancer that interacted ~110 kb away with the promoter of *WNT3* (Fig. 6c), which is known to be involved in cardiac mesoderm induction58,59 and thus may explain the broad cardiac traits potentially affected by this SNP. Because this SNP is within a highly conserved genomic region (Fig. 6d) and predicted to alter a binding motif for the KLF4 transcription factor (Extended Data Fig. 12b), which has been shown to regulate cardiac function in mice^60^, we deleted the putative enhancer containing this SNP in hESCs using CRISPR-mediated genome editing. This deletion consequently resulted in a ~20-50% reduction in *WNT3* expression in mesodermal cells at day 2 (Fig. 6e), but had no effect on expression of the nearest gene *GOSR2* (Extended Data Fig. 12c), which has previously been annotated to rs17608766. Finally, in addition to investigating SNPs associated with cardiac functional traits, we also examined long-range acting SNPs for cardiac developmental traits (i.e. congenital heart disease). Interrogating rs870142, a SNP associated with atrial septal defect^61^, revealed that it was located in an active D2 mesodermal enhancer within the intron of *STX18-AS1*, which formed dynamic long-range interactions (~200 kb) with the promoter of *MSX1* (Fig. 6f, g), a developmental TF reported to interact with *TBX5* during atrial septal development^62,63^. Deletion of this enhancer containing rs870142 using CRISPR/Cas9-mediated genome editing resulted in a 50% reduction in *MSX1* expression (Fig. 6h), whereas the expression of the nearest gene *STX18* remained unchanged (Extended Data Fig. 12d). Thus, these findings support that our chromatin interaction maps can be used to discover candidate genes affected by genetic disease or trait variants located in distal regulatory elements controlling gene expression.

**Figure 6.**
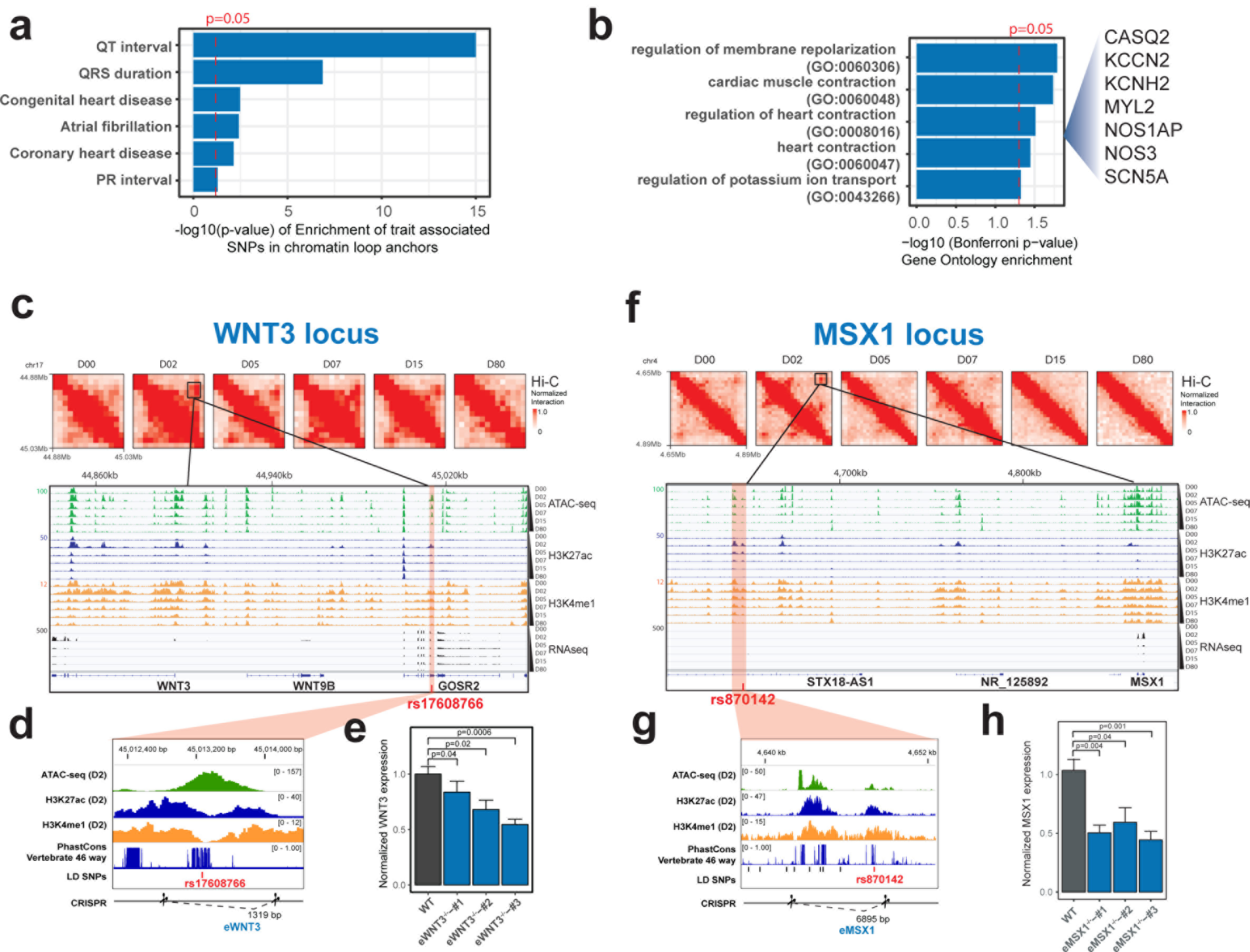
Identification of target gene for heart-related GWAS SNPs using chromatin loops detected during cardiomyocyte differentiation. (**a**) Bar graph shows the enrichment of GWAS SNPs associated with cardiac traits at the chromatin loop anchors detected during the cardiomyocyte differentiation. (**b**) Enriched GO terms for the genes predicted to be linked to SNPs from cardiac GWAS studies along with P values. Red dashed line indicates P value cutoff of 0.05. (**c**) Hi-C contact matrices, and genome browser tracks of various histone modification ChIP-seq and RNA-seq data at the *WNT3* locus. A cardiac GWAS SNP rs17608766 is marked by a red line. (**d**) Expanded view of the genomic region containing rs17608766. The putative enhancer containing rs17608766, which located in an open chromatin region flanked by H3K27ac and H3K4me1 peaks, is targeted for CRISPR/Cas9-mediated deletion. (**e**) Bar charts show the normalized expression levels of *WNT3* in three CRISPR genome edited clones at D2 (mean ± standard error, N =5). Deleting the putative enhancer containing rs17608766 specifically reduces *WNT3* expression. P-values are from one-sided paired t-test. (**f**) Hi-C contact matrices, epigenomic profiles at *MSX1* locus. A cardiac GWAS SNP rs870142 is marked by red line. (**g**) Expanded view of the genomic region containing rs870142. CRISPR mediated deletion of the putative enhancer is marked. (**h**) Bar charts show that deleting the putative enhancer containing rs870142 specifically reduces *MSX1* expression in three independent clones at D2 (mean ± standard error, N = 9). P-values are from one-sided paired t-test.

## DISCUSSION

Evolution of multi-cellular organisms is driven, in large part, by the invention of new gene regulatory circuits responsible for the fitness traits of each species. A long-standing theory holds that retroviruses may play an important role in the evolution of gene regulatory logic^64,65^. Over the years, many classes of endogenous retroviral elements have been found to recruit transcription factors to regulate nearby genes in a cell-type specific manner, or initiate transcription of non-coding RNAs with important regulatory functions^46,66^. Importantly, the expansion of SINE elements in rodents, dogs and opossums have been attributed to the rewiring of gene regulatory networks in these mammals, by expanding the repertoires in each species of CTCF^22^, a DNA binding protein with a critical role in chromatin organization. Surprisingly, no evidence has been found so far for repeat-driven expansion of CTCF binding in the primate genomes^22^, raising the question whether retrotransposon-driven chromatin re-organization indeed is a general strategy of evolution that applies also to the primate lineage. Moreover, retrotransposon elements have been implicated in shaping chromatin boundaries^67^, however, this notion also remains to be experimentally investigated.

Here, we provide multiple lines of evidence to support that the primate-specific retroviral elements HERV-H can delineate TAD boundaries in the human PSCs. Previous studies suggest that HERV-Hs integrated into the human genome during primate evolution to regulate human-specific pluripotency through creating novel chimeric transcripts (*ESRG*^37^, *linc-ROR*^41,68^ and *LINC00458*^49^) and providing potential binding sites to recruit pluripotency factors (*NANOG*, *SOX2* and *POU5F1*)^38^. Our findings reveal that HERV-H sequences might affect gene regulatory program through yet a third mechanism, by shaping the chromatin architecture. Given these HERV-H-mediated chromatin remodeling findings, future investigations of other ERV families of repeats and/or other families of repetitive elements are warranted to illuminate whether other distinct classes of genomic repeats may also exhibit the capacity to shape chromatin architecture in a broad range of cell types including PSCs.

Through correlation analysis and CRISPRi intervention, we reveal that HERV-H’s ability to form TAD boundaries is dependent on its transcriptional activity. In addition, we show that Polymerase II and cohesin complex accumulates at the 3’ terminal of highly transcribing HERV-Hs. Cohesin complex is involved in forming long-range chromatin loops at CTCF binding sites, through an ATP-dependent loop-extrusion mechanism^16,17^. In mammalian cells, most TAD borders are occupied by CTCF and cohesin complex. However, in the human PSC, the HERV-H-associated TAD borders lack canonical CTCF binding motifs and sharp CTCF ChIP-seq peaks, but still retain cohesin complexes. Taken together with the CRISPRi experiment, our findings support that the cohesin complexes are likely positioned by Polymerase II, rather than CTCF. This model is consistent with and provide additional evidence to recent reports of the role of transcription and polymerase movement to position cohesin complex and thus shaping chromatin structure^44,45^.

In addition to the HERV-H discovery, we have interrogated the dynamic chromatin structure re-organization during cardiomyocyte differentiation, cataloged chromatin loops and annotated potential target genes for GWAS variants from cardiac studies. More importantly, we functionally validated our predicted enhancer/SNP to promoter maps by CRISPR mediated deletion, and further showed that WNT3 and MSX1 are likely linked to congenital heart diseases and other cardiac phenotypes. This highlights the utility of the presented chromatin interaction maps to associate non-coding regulatory elements to target genes. Overall, these detailed chromatin accessibility and interaction maps will be a valuable genomic resource for not only providing novel insights into cardiomyocyte development but also identifying important genetic variants associated with cardiac traits including congenital heart disease.

## Acknowledgements

We thank Samantha Kuan and Dr. Bin Li for sequencing and bioinformatics support. We thank Eric van Nostrand for RNA extraction. We would like to thank M. Daadi for providing marmoset iPSCs and F. Gage for providing chimpanzee iPSCs. This project is supported by funding from the Ludwig Institute for Cancer Research (to B.R.), the NIH (1UM1HL128773 to S.M.E., N.C.C., E.D. and B.R.). J.W. is the Virginia Murchison Linthicum Scholar in Medical Research. S.P. was supported by a postdoctoral fellowship from the Deutsche Forschungsgemeinschaft (DFG, PR 1668/1-1). E.N.F. was supported by NIH pre-doctoral training grant (5T32HL007444-35).

## Author contributions

N.C.C. and B.R. designed and supervised the experiments, analysis, and data interpretation. Y.Z. implemented the analysis pipeline and analyzed all sequencing datasets, interpreted the results, designed the experiments for HERV-H and chromatin loop functional studies. T.L. generated the CRISPR-cas9 edited cell lines for HERV-H and chromatin loop functional studies, performed differentiation and qPCR of the corresponding cell lines. S.P. initiated the project, performed the Hi-C and ATAC-seq experiments, and helped with interpretation of the results. J.G. and E.N.F performed cell culture, differentiation and collected cells for Hi-C, ChIP-seq, ATAC-seq and RNA-seq assays. E.D. contributed to analysis and interpretation of the ChIP-seq data. ChIP-seq experiments were performed by A.Y.L. (histone modifications), S.C. (CTCF), Q.Z. and H.H. (SMC3). Y.Q. and R.F. helped with the analysis of Hi-C datasets. K.M. helped with the genome-editing experiments. L. Y., J. C. I. B and J. W. cultured and prepared non-human primate iPSCs for sequencing. Z.Y. performed the RNA-seq experiments. R.H. performed the Hi-C experiments for HERV-H knock-out, CRISPRi, HERV-H knock-in and primate iPSC cell lines. S.M.E. helped with interpretation of the results. Y.Z., T.L., S.P., N.C.C., and B.R. wrote the manuscript with input from all authors.

## Author Information

Sequencing data from this study have been deposited in the Gene Expression Omnibus under the accession number GSE116862. The authors declare no competing financial interests. Readers are welcome to comment on the online version of the paper. Correspondence and requests for materials should be addressed to N.C.C. and B.R.

## Extended Data Figure Legends

**Extended Data Figure 1.**
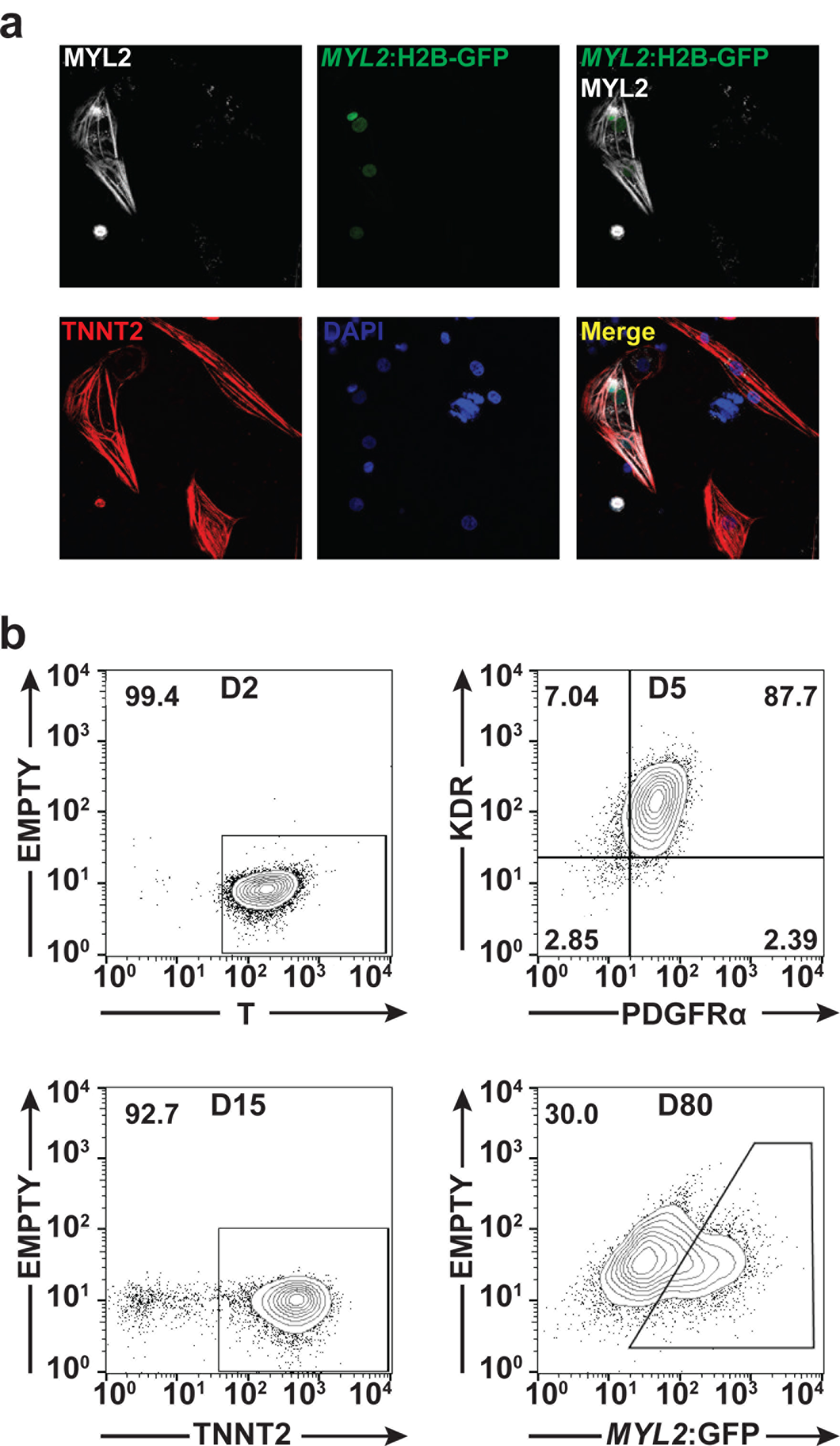
Imaging and Flow cytometry data for cardiomyocyte differentiation. (**a**) Immunostaining for MYL2 (white) shows exclusive expression in *MYL2*-H2B-GFP + hESC-derived cardiomyocytes after 80 days of cardiac differentiation. *MYL2*-H2B-GFP + cells were positive for cardiac troponin T (cTNT) (red), but not all cTnT+ cells were MYL2+ cardiomyocytes. DNA was stained with DAPI (blue). Images are representative of a minimum of three independent experiments. (**b**) Flow cytometric quantification of distinct cell-surface markers for D2-D15 time points: T (D2), KDR/PDGFRα (D5), TNNT2 (D15). H2B-GFP + ventricular cardiomyocytes were sorted at day 80. Numbers represent the respective percentage of cells.

**Extended Data Figure 2.**
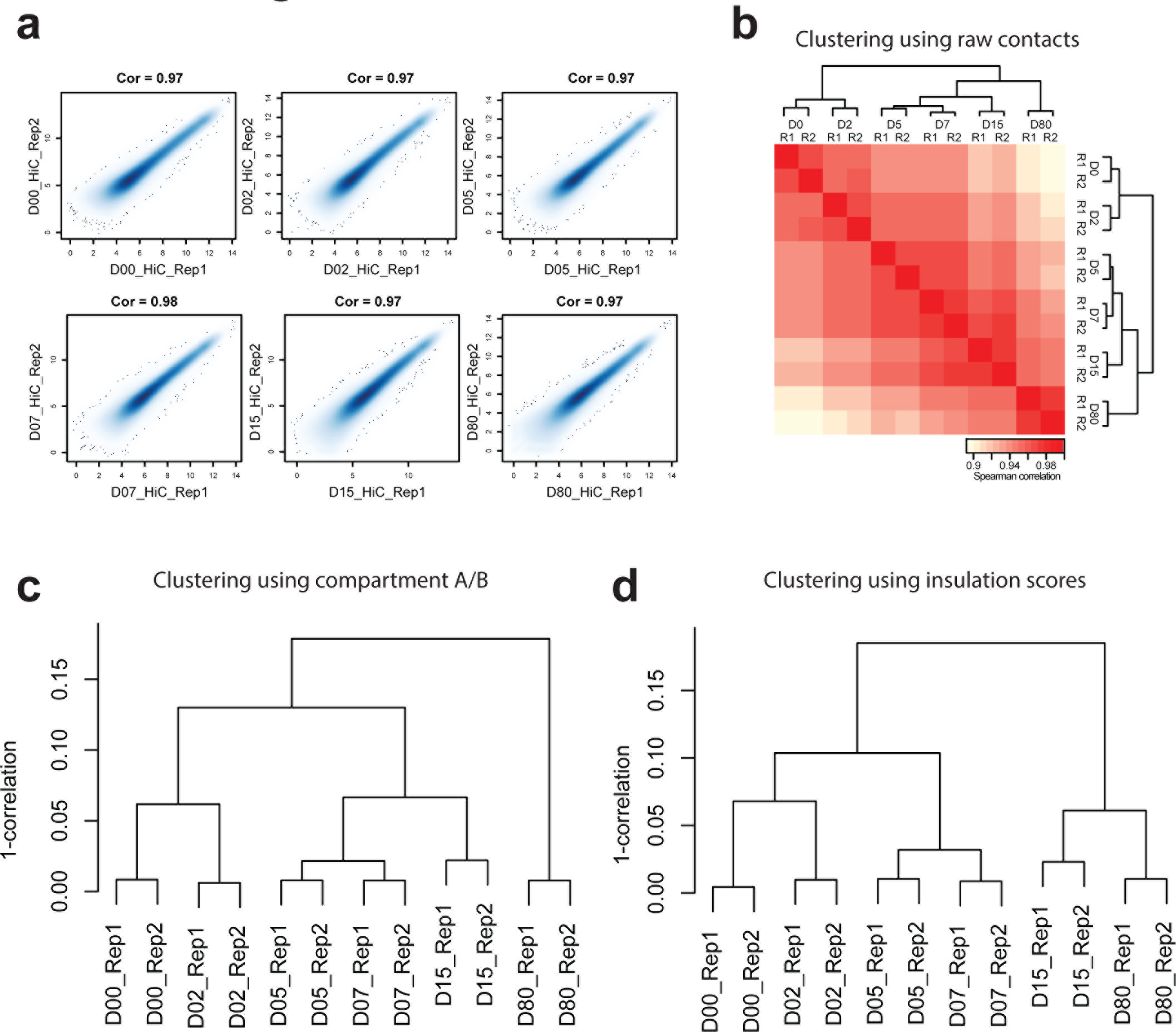
The Hi-C data are highly reproducible. (**a**) Smoothed scatterplot of the Hi-C contacts between biological replicates for each stage. bin size = 100 kb. (**b**) Hierarchical clustering of Hi-C contact matrices based on 1-PCC (Pearson correlation coefficients) of the contacts between all samples, bin size = 100 kb. (**c**) Hierarchical clustering of the compartment A/B scores (PC1 values) between samples. (**d**) Hierarchical clustering of the insulation scores between samples.

**Extended Data Figure 3.**
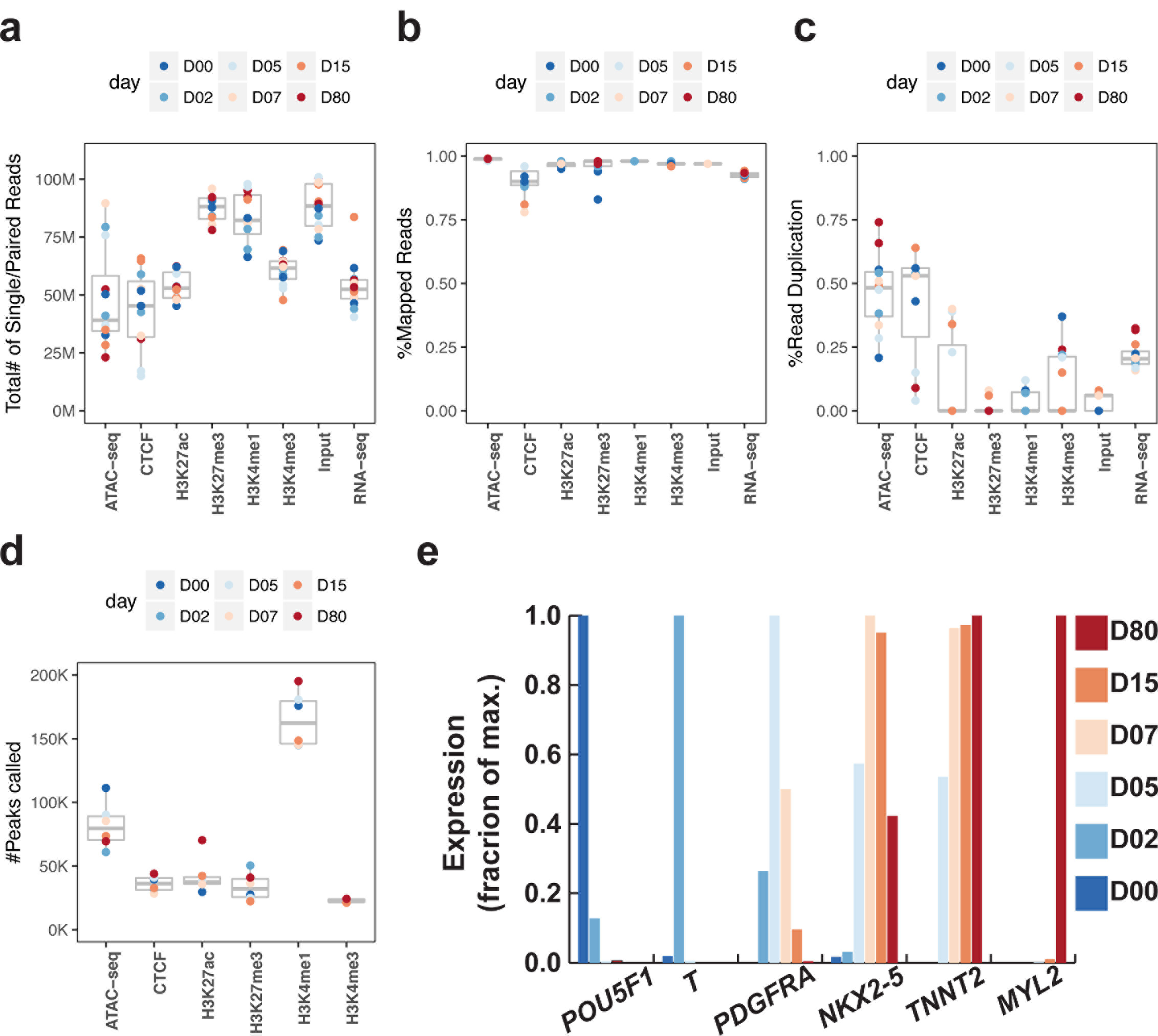
Quality metrics of ChIP-seq, RNA-seq and ATAC-seq data used in the present study. (**a**) Boxplot showing the number of reads for each sample. (**b**) Boxplot showing the percentage of alignments to human genome for each sample. (**c**) Boxplot showing the percentage of potential duplicated reads for each sample. (**d**) Boxplot showing the number of peaks called for each ChIP-seq and ATAC-seq sample. (**e**) Barplots showing the relative expression levels (by RNA-seq) of representative genes during cardiomyocyte differentiation.

**Extended Data Figure 4.**
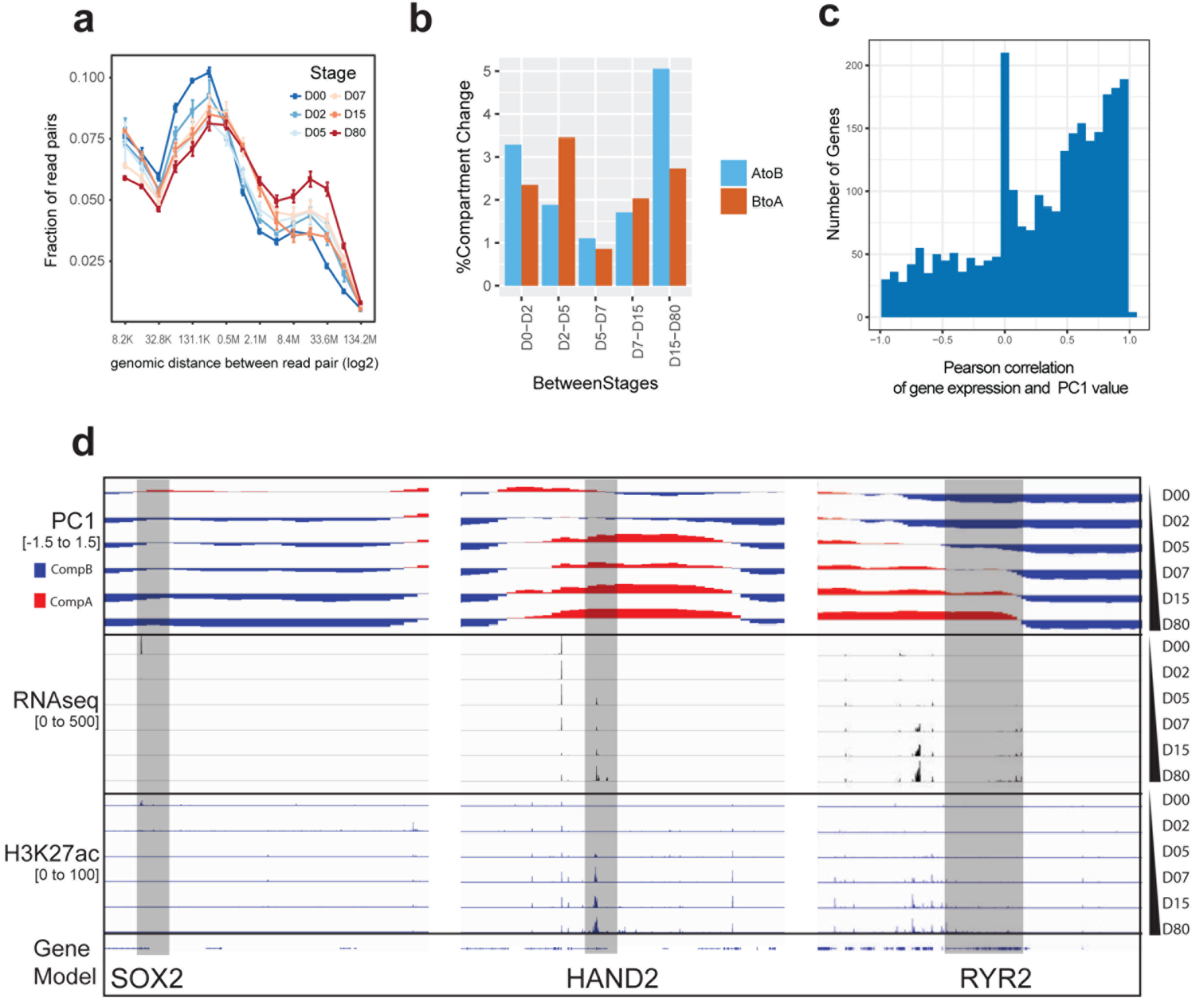
Global changes in chromatin organization during cardiomyocyte differentiation. (**a**) Probability of Hi-C interaction versus interacting distance (log2) during cardiomyocyte differentiation. (**b**) Barplot shows the percentages of compartment switches for each stage transition. (**c**) Histogram of the Pearson correlation coefficients between gene expression and compartment A/B (PC1) value. (**d**) Genome browser view of *SOX2*, *HAND2* and *RYR2* loci, showing PC1 (Blue/red), RNA-seq (Black) and H3K27ac (Blue) signals.

**Extended Data Figure 5.**
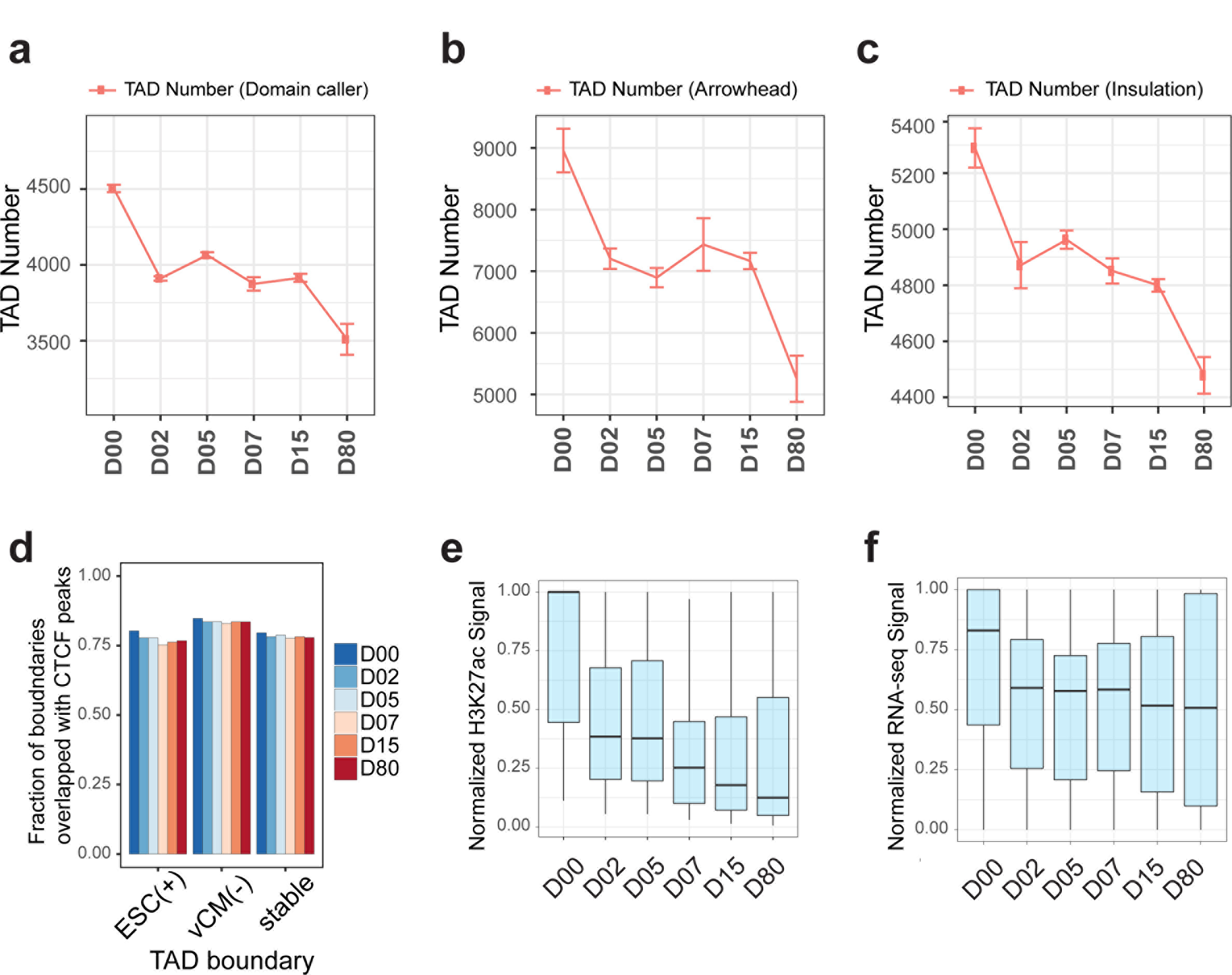
Dynamics of TADs during cardiomyocyte differentiation. (**a**) Number of TADs as defined by a domain caller algorithm first reported by Dixon et al. 2012^5^. (**b**) Number of TADs (or loop domains) found by Arrowhead algorithm^14^. (**c**) The number of TADs found by Insulation score^34^. (**d**) Fraction of TAD boundaries that contain CTCF ChIP-seq peaks at each stage of cardiomyocyte differentiation. “stable” group stands for TADs that are present at all stages. (**e**) Boxplot showing the dynamics of H3K27ac signal at ESC(+) TADs. (**f**) Boxplot showing the dynamics of RNA-seq signal at ESC(+) TADs.

**Extended Data Figure 6.**
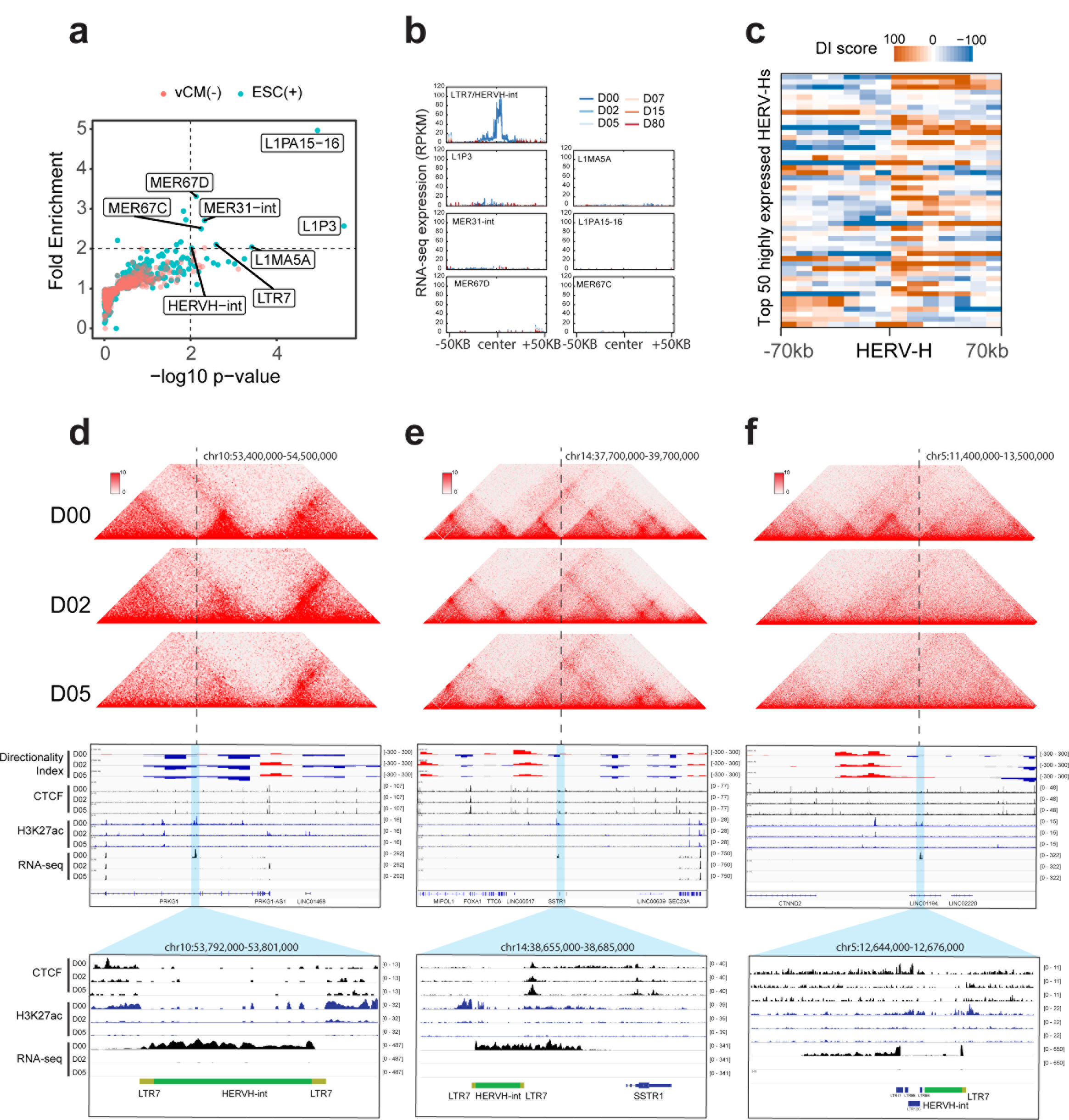
HERV-H elements are enriched at ESC(+) TAD boundaries. **(a)** Scatter plot shows the fold enrichment and −log10(p-values) for various repeat element classes at ESC(+) and vCM(-) TAD boundaries relative to the rest of the TAD boundaries. P-values are from two-sided proportion test. (**b**) Aggregate RNA-seq expression profile (RPKM normalized) at ESC(+) TAD boundaries that overlap each of the eight repeat elements (HERVH-int and LTR7 are combined as they both belong to HERV-H). (**c**) Heatmap of DI scores within 70 kb of the top 50 most highly-expressed HERV-Hs. (**d,e,f**) Hi-C heatmap, DI, and epigenomic profiles for three ESC(+) TAD boundaries harboring HERV-H/LTR7 sequences.

**Extended Data Figure 7.**
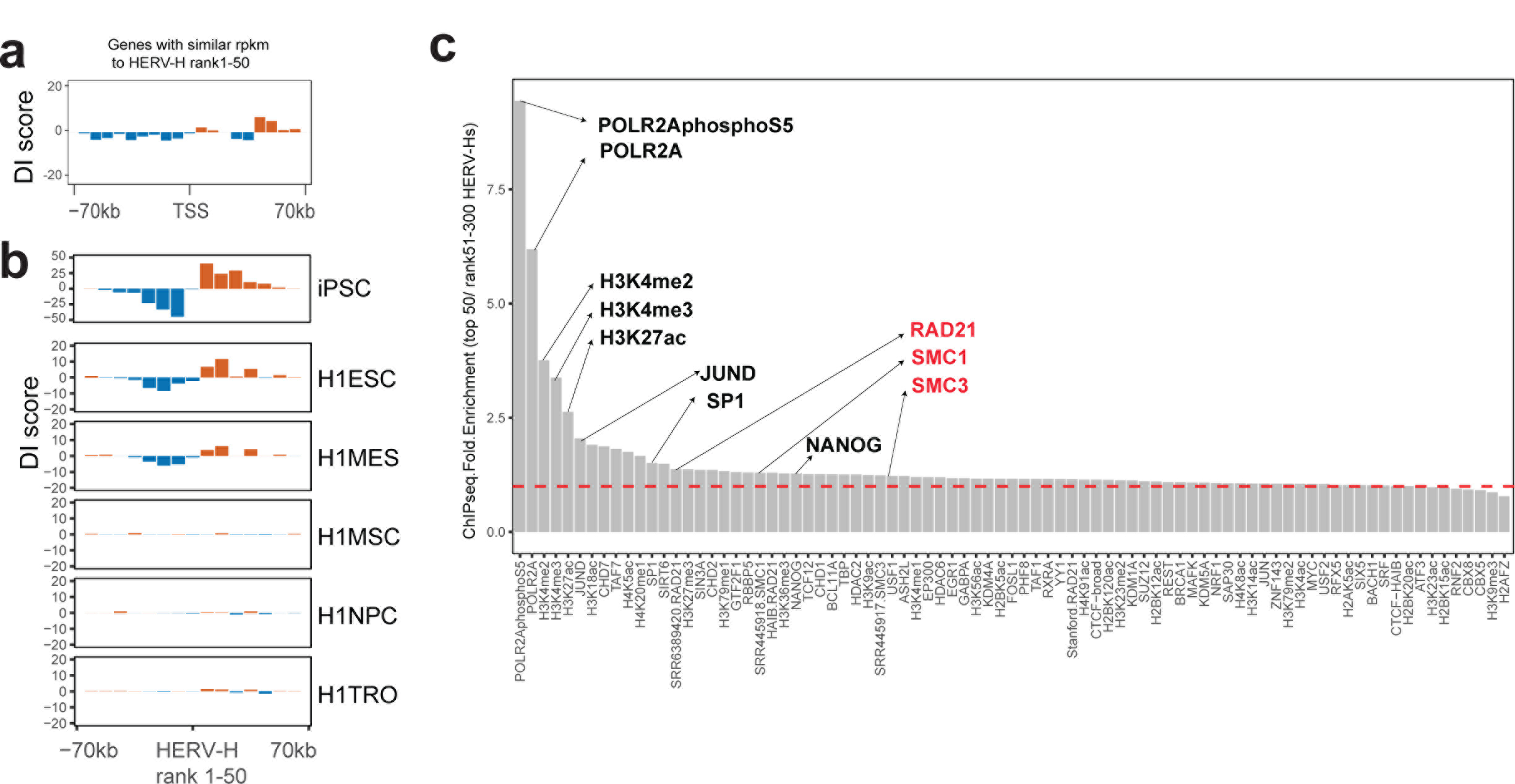
Further characterization of the HERV-H-associated TAD boundaries. (**a**) DI score profiles around the TSSs of genes expressed at similar levels to the top 50 transcribed HERV-Hs. (**b**) Aggregated DI score profiles of the TAD-boundary associated with HERV-Hs in multiple H1 ESC derived lineages and iPSCs. ESC: embryonic stem cell; MES: mesendoderm. MSC: mesenchymal stem cell; NPC: neural stem cell; TRO: trophoblast-like cell; iPSC: induced pluripotent stem cell. Interestingly, in human mesendoderm cells (an early human embryonic state that gives rise to mesoderm and endoderm cells), both the expression of these HERV-Hs and their corresponding TAD boundary strength were approximately half the levels compared to those observed in hESCs. There might be some un-differentiated cells in this population. (**c**) Barplot shows the ChIP-seq signal fold enrichment of the top 50 HERV-Hs comparing to HERV-Hs ranking 51-300.

**Extended Data Figure 8.**
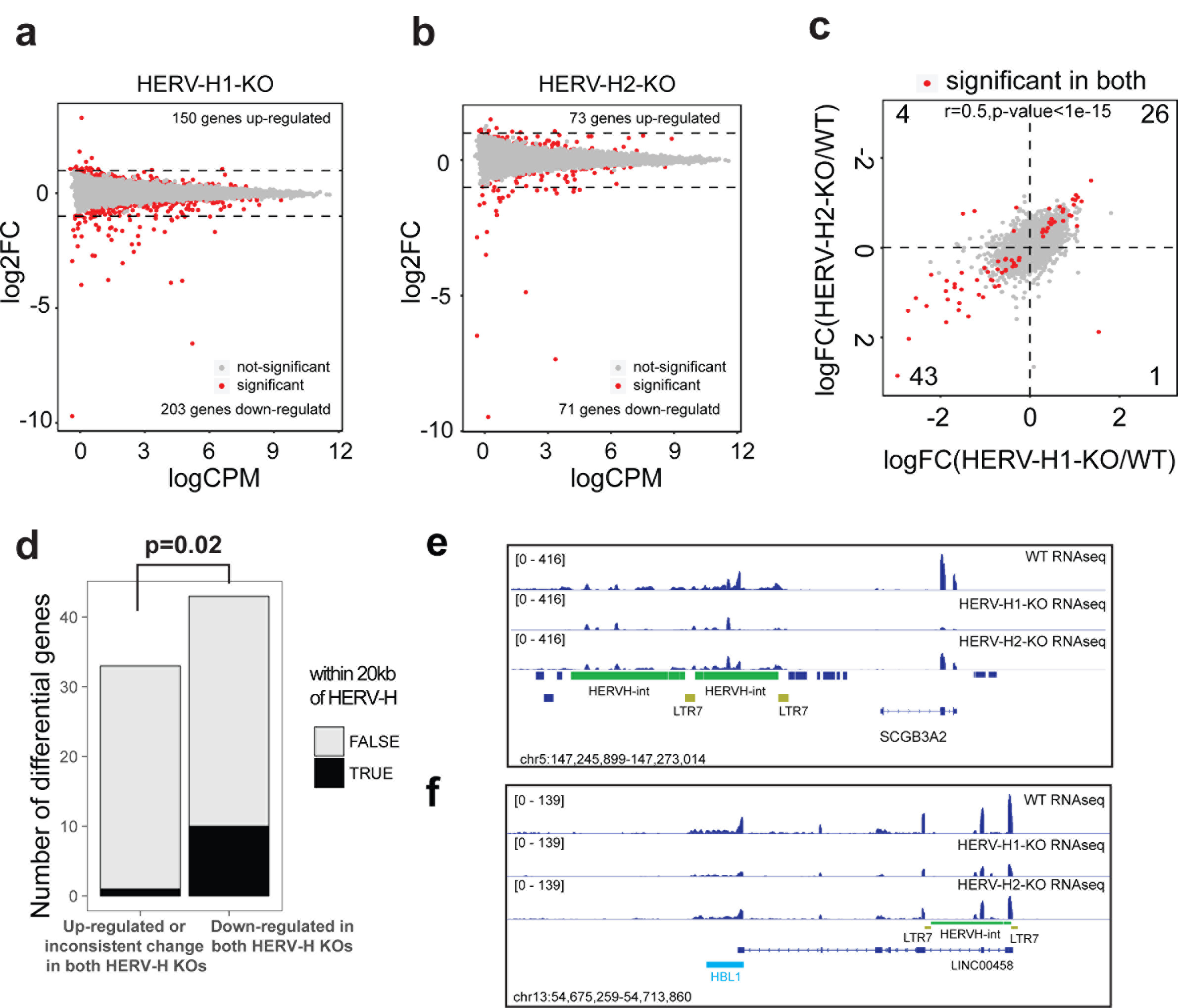
Changes in gene expression in HERV-H knockout hESCs. (**a**) MA plot showing average gene expression levels and fold changes of each gene in HERV-H1-KO and wild-type (WT). (**b**) Same as (**a**) but for HERV-H2-KO. (**c**) Scatterplot shows the changes in gene expression in HERV-H1-KO and HERV-H2-KO cells over WT cells. The red dots are genes significantly changed in both mutant cell lines. The numbers at the corners show the number of significantly changed genes in each Quadrant. (**d**) Barplot showing the number of significantly changed genes located within 20 kb of the HERV-H sequences. Genes down-regulated in both HERV-H knockouts were more likely to be within 20 kb of HERV-H sequences. (**e**, **f**) RNA-seq profile of wild-type (*WT*) and HERV-H1-KO and HERV-H2-KO lines at the *SCGB3A2* and *LINC00458/HBL1* gene loci.

**Extended Data Figure 9.**
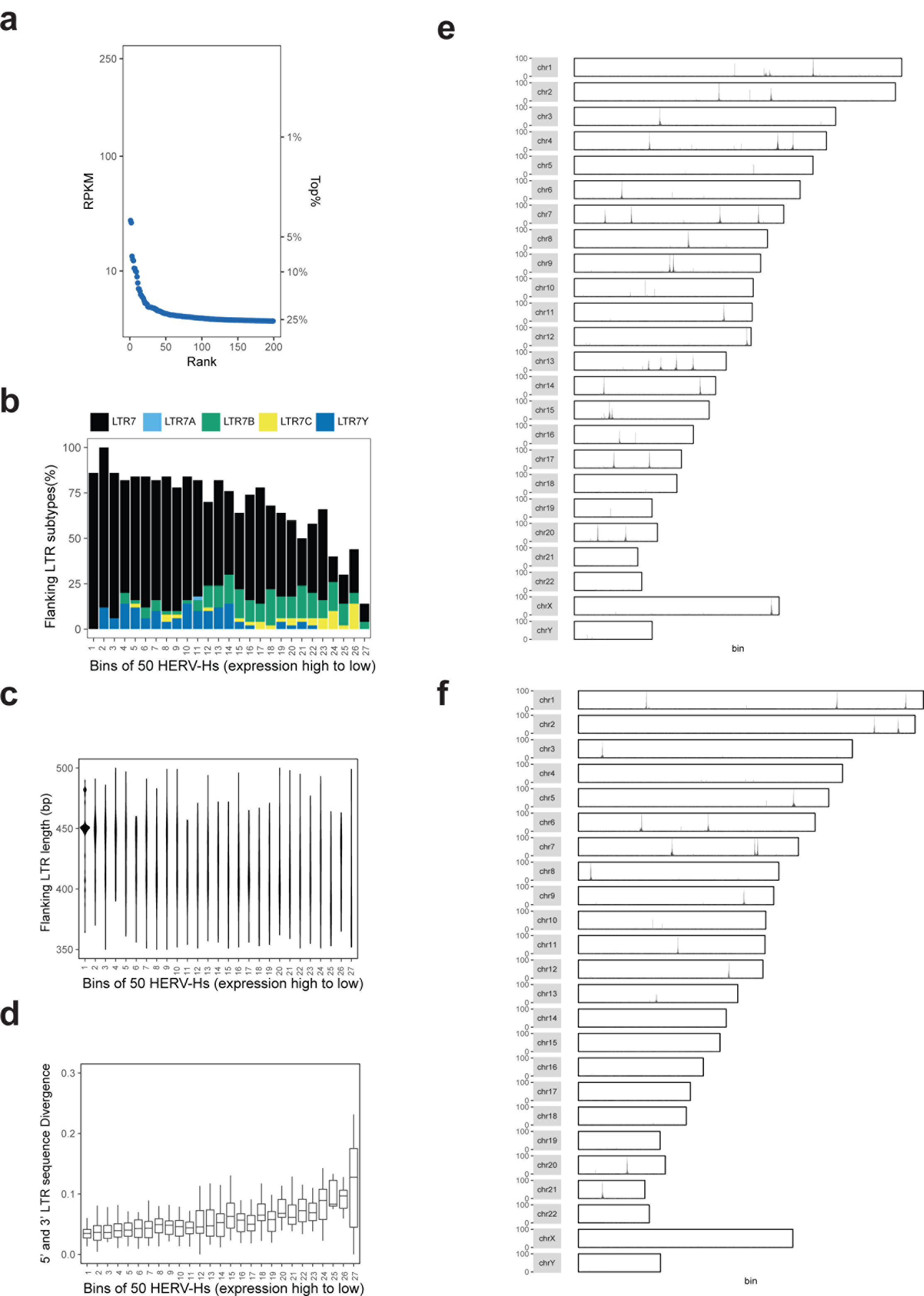
Analysis of HERV-H LTR sequences and HERV-H insertion. (**a**) Scatterplot shows the expression levels (RPKM) across different HERV-H loci in chimpanzee iPSC (ordered by expression levels from high to low). The expression of HERV-Hs is at least 10 fold less than their human counterparts. (**b**) Bar graph shows the percentage of HERV-Hs flanked by various types of LTRs. HERV-Hs are ranked by their expression levels in the hESCs, and grouped by bins of 50. (**c**) Violin plot shows the length of the flanking LTRs. HERV-Hs ranked and binned as described in (**b**). (**d**) Boxplot shows the sequence divergence of the 5’ LTR and 3’ LTR for each bin of HERV-Hs. HERV-Hs ranked and binned same as (**b**). (**e**) The insertional profile of HERV-H in the HERV-H-ins.clone1 mutant line. The y-axis shows the pileup of Hi-C read pairs with one end mapped to the HERV-H2 sequence. Based on proximity ligation principle, loci with high pileup should harbor HERV-H2 insertion. (**f**) Same as (**e**) but for HERV-H-ins.clone2 mutant line.

**Extended Data Figure 10.**
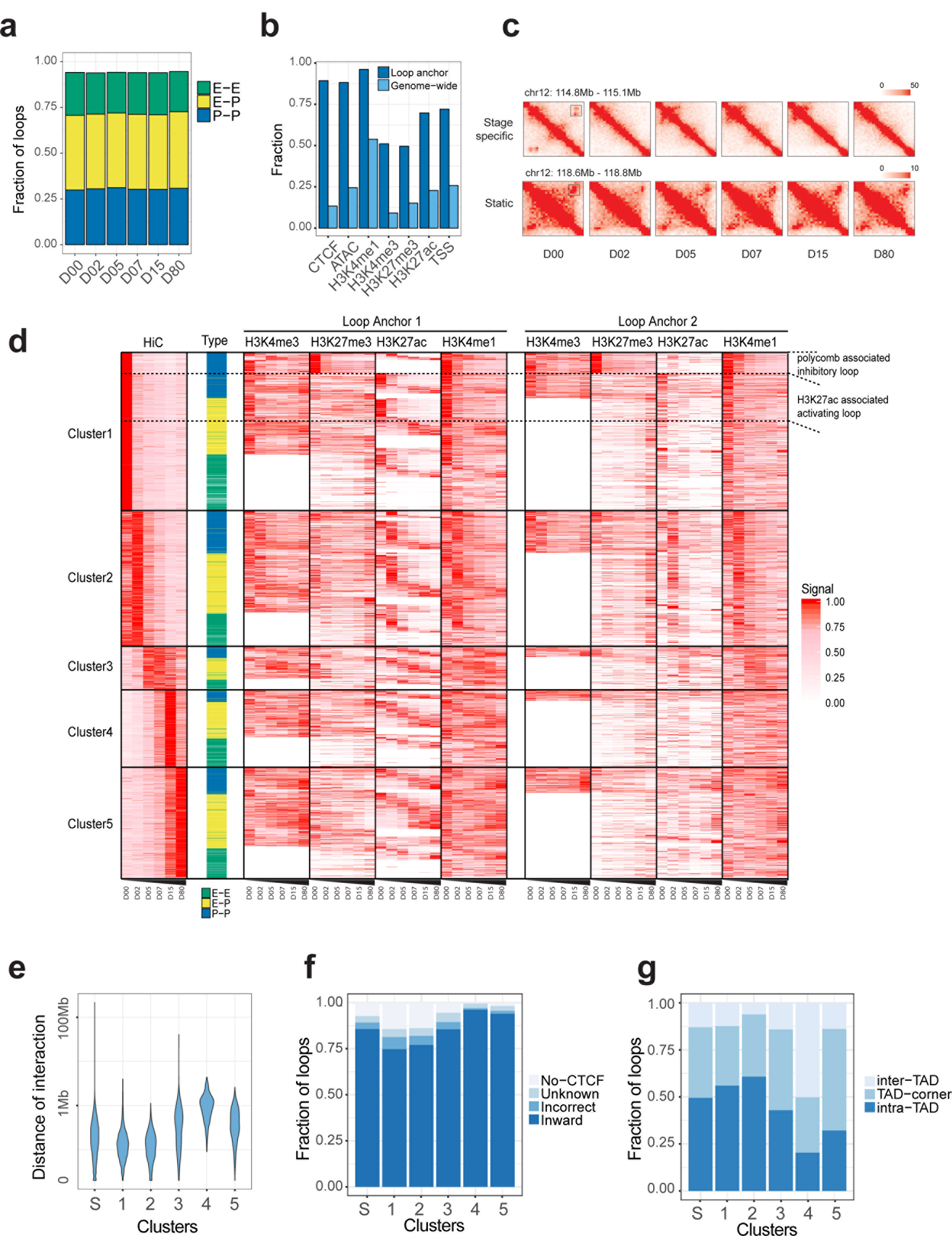
Relationships between chromatin loops and histone modification during cardiomyocyte differentiation. (**a**) Barplot showing fraction of chromatin loops in each category of interactions. E-E: enhancer to enhancer; E-P: enhancer to promoter; P-P: promoter to promoter. (**b**) Barplot showing fraction of chromatin loop anchors or genome-wide background containing each of the epigenomic element. (**c**) Representative data for Hi-C contact matrices of stage-specific and static chromatin loop interactions (**d**) Heatmap shows the clustering of stage-specific chromatin loops and histone modification states at the corresponding loop anchors. (**e**) Violin plot of the genomic distances for each class of chromatin loops in log scale. Symbol S: static loop. 1-5: Loop clusters 1-5 in **Fig.4b** and Ext. Fig. 10d. (**f**) Barplot for fraction of loops harboring different type of CTCF orientations. No-CTCF: at least one anchor does not contain CTCF peak; unknown: at least one anchor does not contain a matched CTCF motif; incorrect: CTCF motifs at both anchors do not face inward; inward: CTCF motifs at both anchors face inward. (**g**) Barplot for fraction of loops in each category related to TAD boundaries. Inter-TAD: loop is outside of all TADs; TAD-corner: loop is within 20 kb of any TAD corner; intra-TAD: loop is inside of TAD.

**Extended Data Figure 11.**
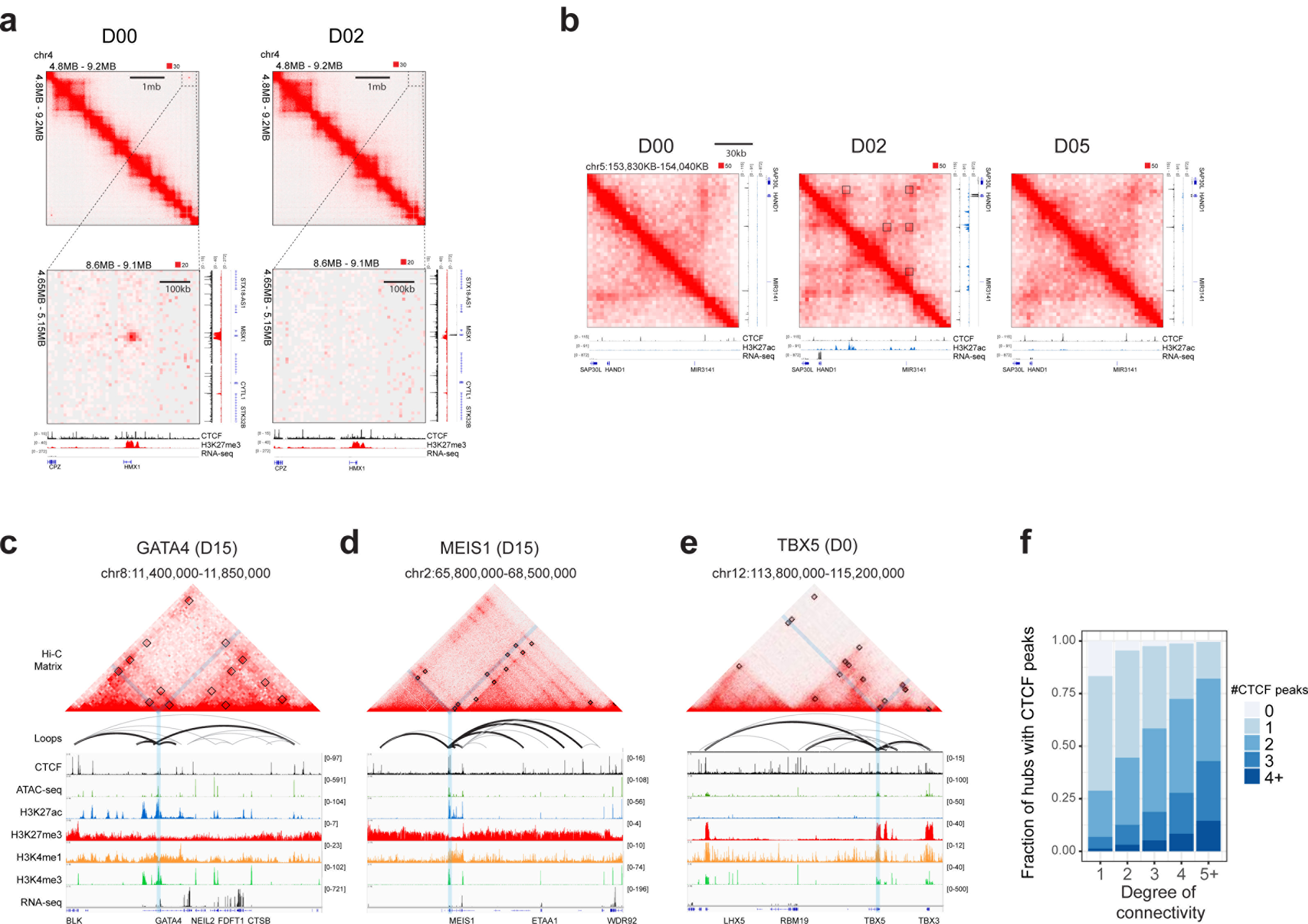
Chromatin interaction hubs around genes encoding cardiac transcription factors. (**a**) Example of a H3K27me3-associated inhibitory loop connecting *MSX1* and *HMX1* promoters. (**b**) Example of several stage-specific and H3K27ac associated activating loops at *HAND1* locus. (**c,d,e**) Hi-C contact matrices (top), loop arcs (middle), and epigenomic profiles (bottom) for *GATA4* (**c**), *MEIS1* (**d**) and *TBX5* (**e**). (**f**) Fraction of network hubs containing CTCF peaks plotted against the degree of connectivity as illustrated by the number of CTCF peaks at each anchor connectivity.

**Extended Data Figure 12.**
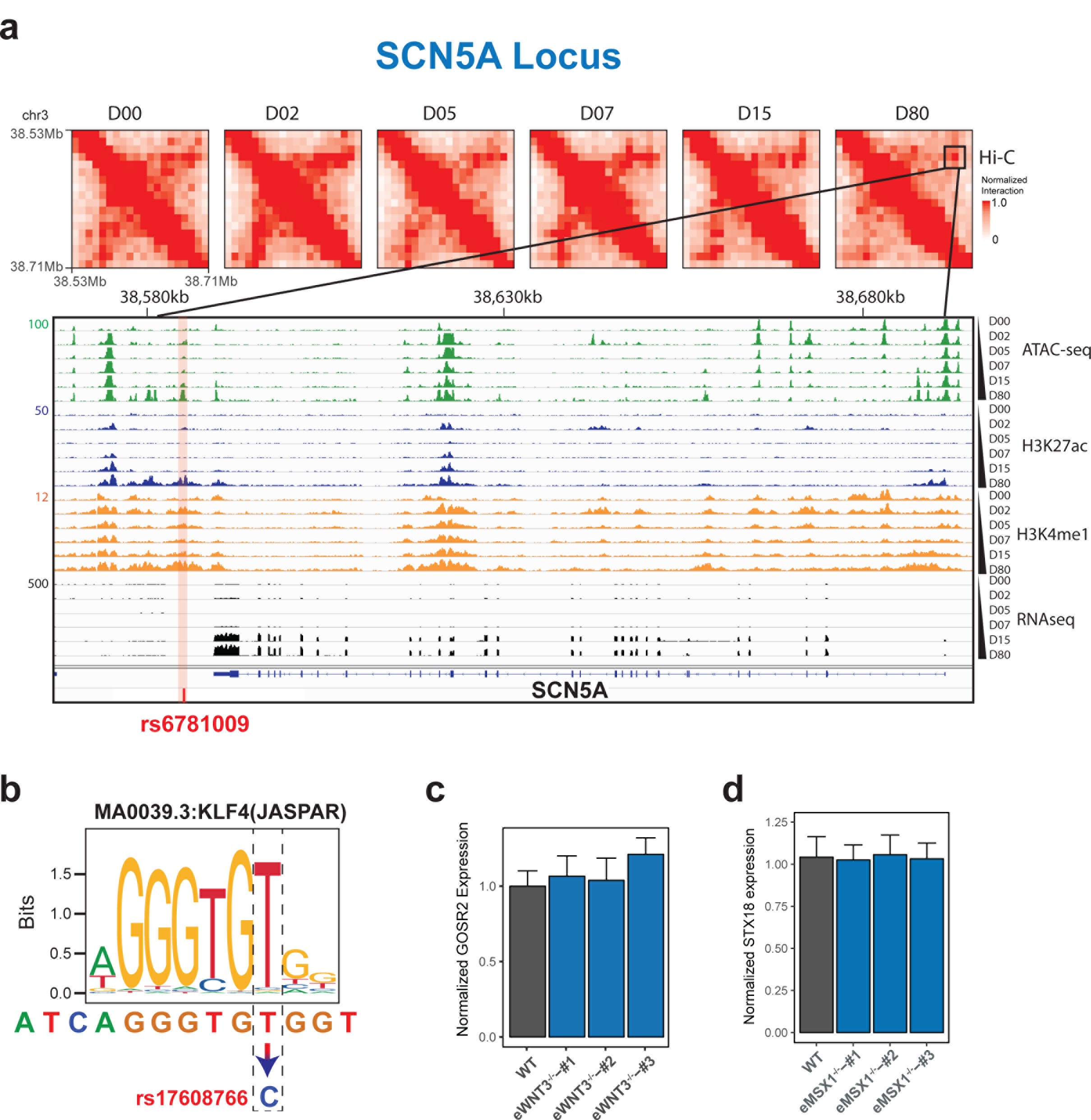
Validation of target gene predictions for heart-related GWAS SNPs. (**a**) Hi-C contact matrices, epigenomic profiles (ATAC-seq, H3K4me1, H3K27ac and RNA-seq) and cardiac GWAS SNP rs6781009 at *SCN5A* locus. (**b**) rs17608766 is predicted to alter KLF4 motif in the putative enhancer. (**c**) Normalized expression values of *GOSR2* in three CRISPR-genome edited clones shows that deleting the putative enhancer containing rs17608766 does not change *GOSR2* expression (mean ± standard error, N=5). (**d**) Normalized expression values of *STX18* in three CRISPR-genome edited clones shows that deleting the putative enhancer containing rs870142 does not change *STX18* expression (mean ± standard error, N=7).

**Extended Data Table 1.** Summary statistics for Hi-C data

**Extended Data Table 2.** List of stage-specific TAD boundaries.

**Extended Data Table 3.** List of differentially expressed genes in HERV-H1-KO and HERV-H2-KO.

**Extended Data Table 4.** List of chromatin loops.

**Extended Data Table 5.** List of genes located on the network hubs.

**Extended Data Table 6.** List of target genes assigned to cardiac GWAS SNPs with chromatin loops.

**Extended Data Table 7.** List of antibodies used for ChIP-seq.

**Extended Data Table 8.** List of primers and cell lines used in this study.

